# Genome-wide annotation of gene regulatory elements linked to cell fitness

**DOI:** 10.1101/2021.03.08.434470

**Authors:** Maria ter Weele, Tyler S. Klann, Alejandro Barrera, Adarsh R. Ettyreddy, Siyan Liu, Ryan A. Rickels, Julien Bryois, Simon Jiang, Shaunak S. Adkar, Nahid Iglesias, Patrick F. Sullivan, Timothy E. Reddy, Andrew S. Allen, Gregory E. Crawford, Charles A. Gersbach

## Abstract

Noncoding regulatory elements control gene expression and thus govern nearly all biological processes. Epigenomic profiling assays have identified millions of putative regulatory elements, but systematically determining the function of those regulatory elements remains a substantial challenge. Here we adapt CRISPR screening by epigenetic repression to screen all 111,619 putative non-coding regulatory elements defined by open chromatin sites in human K562 leukemia cells for their role in regulating essential cellular processes and proliferation. In an initial screen containing 1,084,704 gRNAs, we implemented an analysis framework to quantify perturbation effects, and nominate 1,108 regulatory elements that strongly impact cell fitness. We tested 8,845 of the primary screen elements in a secondary screen, evaluated their cell-type specificity in a second cancer cell line, and then used a single-cell RNA-seq CRISPR screen to discover 63 connections between distal regulatory elements and target genes. This comprehensive and quantitative genome-wide map of essential gene regulatory elements presents a framework for extensive characterization of noncoding regulatory elements that drive complex cell phenotypes and for prioritizing non-coding genetic variants that may contribute to common traits and disease risk.

## Introduction

Human gene regulatory elements control gene expression and orchestrate many biological processes including cell differentiation (Nguyen et al. 2015), proliferation (Sur and Taipale 2016), and environmental responses (Ghisletti et al. 2010). Genetic and epigenetic variation that alters gene regulatory element function is a primary contributor to human traits and susceptibility to common disease (Maurano et al. 2012). Studies of chromatin state and transcription factor occupancy have identified millions of putative human gene regulatory elements (Thurman et al. 2012). The biological importance and large number of putative human gene regulatory elements have motivated the development of high-throughput technologies to measure regulatory element activity genome-wide. Examples include assays that measure the influence of putative regulatory elements on reporter gene expression (Arnold et al. 2013; Johnson et al. 2018), and targeted CRISPR-based methods to measure the effects of genetic or epigenetic perturbation of up to thousands of regulatory elements in their native chromosomal context (Montalbano, Canver, and Sanjana 2017).

A key measure of gene or regulatory element function is its contribution to overall cell fitness, comprising the balance of cell survival and proliferation. Genome-wide perturbation technologies, such as RNAi and CRISPR-based screens, have identified genes and noncoding RNAs involved in diverse essential cellular processes (Rauscher et al. 2017; Lenoir, Lim, and Hart 2018; E. E. Schmidt et al. 2013; Hart et al. 2015; Wang et al. 2017, 2014; Shalem et al. 2014; Haswell et al. 2021; Raffeiner et al. 2020). CRISPR-based genetic or epigenetic perturbation of noncoding regulatory elements within specific genomic loci have identified target genes and downstream effects on cell phenotypes (Canver et al. 2015; Rajagopal et al. 2016; Fulco et al. 2016; Sanjana et al. 2016; Klann et al. 2017; Gasperini et al. 2018; P. B. Chen et al. 2022). However, these perturbation screens of distal regulatory elements have generally been limited to targeted regions of the genome or loci encoding oncogenes (Klann, Black, and Gersbach 2018). Consequently, functional understanding of the millions of predicted human gene regulatory elements remains sparse, making it difficult to routinely establish gene regulatory contributions to human traits and disease.

Here, we identify >1,000 human gene regulatory elements that functionally contribute to cell fitness. We used a genome-wide CRISPR-based repression screen that individually targeted each of the >100,000 putative gene regulatory elements in the human K562 erythroleukemia cell line. We further characterize the properties, distribution, cell-type specificity, and target genes of the identified regulatory elements. These elements and target genes confirm and complement results from gene-based screens, and suggest new pathways and molecular processes that contribute to cell fitness. Comprehensive annotations of regulatory element function, such as the study presented here, are critical for creating new reference datasets to prioritize regulatory elements and noncoding variants that contribute to human traits and diseases.

## Results

### Genome-wide screen of all regulatory elements in K562 cells

We used a whole-genome CRISPR repression (CRISPRi) screen to measure the effect of epigenetically silencing 111,619 unique putative regulatory elements, defined by DNase-I hypersensitive sites (DHS), on cell fitness in K562 cells (**Fig. 1A**, **Fig. S1**) (ENCODE Project Consortium 2012). We assayed K562 cells because they are suspension cells and one of the most extensively characterized cell models in terms of chromatin accessibility, histone marks, transcription factor binding, and gene expression (ENCODE Project Consortium et al. 2020). Our library contained 1,084,704 unique gRNAs averaging ∼10 gRNAs per DHS (**Fig. S2A, Table S1**), and we transduced this library into a clonal K562 cell line stably expressing the dCas9^KRAB^ transcriptional repressor (**Fig. 1B**) (Gilbert et al. 2013; Thakore et al. 2015). After ∼14 population doublings, we quantified gRNA abundance across 3 experimental and 4 control replicates with robust count correlations (Pearson correlation > 0.9, **Fig. S2B**). Depletion induced by dCas9^KRAB^ in comparison to unmodified cells indicates that the targeted regions contribute to cell fitness. As this was a discovery screen, we used a relaxed false discovery rate (FDR) of 0.1, and identified a significant depletion of 13,732 gRNAs in 11,186 DHSs (FDR < 0.1, **Fig. 1C**), largely with 1 significant gRNA per DHS (**Fig. S2C**). To avoid discarding signals below an arbitrary gRNA significance cutoff, we implemented a stringent FDR-controlled aggregation of all gRNAs across the DHS, using non-essential gene promoter DHSs as empirical controls (see **Methods**, **Fig. S3A**) (Hart et al. 2014). We identified 1,038 high-confidence significantly depleted DHSs (FDR < 0.05), indicating that repressing those DHSs impaired cell viability or proliferation (**Fig. 1D-E**). We also found 4,196 significantly enriched gRNAs across 4,025 DHSs (FDR < 0.1, **Fig. 1C**), indicative of perturbations to gene expression that increased cell fitness. Using the same aggregation method, only 70 high-confidence DHSs were significantly enriched (FDR < 0.05), representing elements that can be repressed to enhance cell fitness (**Fig. 1D-E, Table S2**). After this stringent aggregation, most significant DHSs contained multiple individually significant gRNAs, though a smaller number contained only 1 or no individually significant gRNAs (**Fig. S3B**).

**Figure 1:**
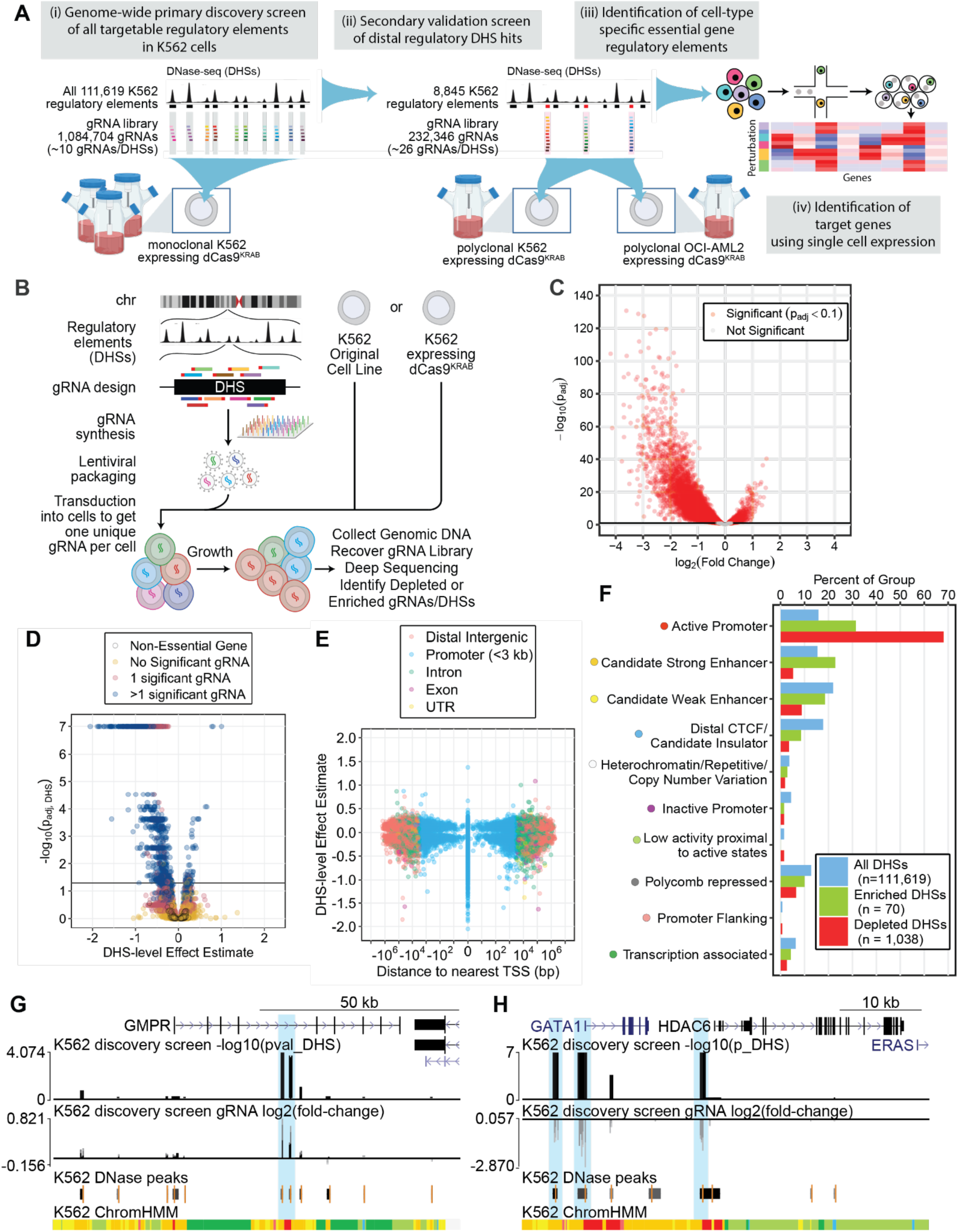
Whole-genome CRISPRi screen identifies regulatory elements essential for K562 cell fitness. **(A)** Overall schematic of (i) discovery screen, (ii) secondary validation screen of regulatory elements, (iii) cell-type specificity analysis, and (iv) single-cell RNA-seq readout to connect cell fitness-associated regulatory elements to target genes. **(B)** A schematic of the approach to screening cell fitness. gRNAs are designed to target all DHSs in the K562 cell line and synthesized as a pool for lentiviral delivery. K562 cells either constitutively expressing or not expressing dCas9^KRAB^ are treated with the lentiviral gRNA library at a low MOI and cultured for 14 population doublings. Genomic DNA is harvested and gRNA abundance is quantified by Illumina sequencing. **(C)** Significance of gRNA abundance changes relative to log_2_(fold-change). **(D)** Significance of DHS effect on cell growth relative to aggregate effect size estimate, colored by the number of individually significant gRNAs in the DHS (FDR < 0.1). Non-essential gene promoters, used as negative controls in the aggregation, are separately indicated. A DHS-level false discovery rate (FDR) of 0.05 is marked with a horizontal line. **(E)** Distribution of DHS-level effect estimates relative to distance to the nearest transcriptional start site. **(F)** Relative abundances of DHSs across chromHMM annotation classes, including the full set, significantly enriched DHSs, and significantly depleted DHSs. **(G)** Representative genome browser tracks of enriched DHSs in the intron of GMPR (hg19 chr6:16,208,540-16,310,537). Highlighted regions are statistically significant (FDR < 0.05). **(H)** Representative genome browser tracks of depleted DHSs near GATA1. Highlighted regions are statistically significant (FDR < 0.05).

### Attributes of gRNAs that impact cell fitness

Effect sizes for gRNAs that reduced cell fitness were overall greater (average log_2_(fold-change) = −0.67) than those that increased cell fitness (average log_2_(fold-change) = 0.259, **Fig. 1C**). This is consistent with it being easier to reduce fitness than increase fitness of the rapidly growing K562 cell line in its standard culture media. To better understand the characteristics that distinguish the significantly enriched or depleted gRNAs, we annotated each gRNA in the library with a selection of features (**Fig. S4**). In the non-essential negative control DHSs, there were no trends between gRNA sequence features and enrichment. More globally across all DHSs, there are slight correlations between protospacer nucleic acid composition and significance in the discovery screen. The correlation between GC content and gRNA significance is stronger in essential DHSs: within significant DHSs, high GC content and low T content is associated with more effective gRNAs. The association is not simply driven by the “TTTT” U6 termination motif known to be associated with poor gRNA performance in similar vectors, as none of the gRNAs analyzed contained the motif (Yao et al. 2024). Notably, there was no correlation between gRNA significance in the genome-wide discovery screen and cutting frequency determination (CFD) scores (**Fig. S4B**), a gRNA specificity metric (H. Schmidt et al. 2022; Perez et al. 2017). Whereas gRNAs with CFD scores less than 0.2 were previously reported to associate with a negative impact on cell fitness due to low-specificity and toxicity (Tycko et al. 2019), this does not appear to be driving signal in this screen.

### Attributes of DHSs that impact cell fitness

While significant DHS perturbations were found at a range of distances from their nearest gene (median absolute distance 12.25 kb, IQR 2.43 to 37.59 kb), the strongest observed signals centered on DHSs that overlapped with FANTOM transcriptional start sites (TSSs, **Fig. 1E**) (Horlbeck et al. 2016). This is consistent with previous studies showing that repressing promoters with dCas9^KRAB^ has a larger effect on gene expression than repressing distal regulatory elements (Thakore et al. 2015; Klann et al. 2017). Although overall effect sizes and significance levels decrease with distance from TSSs, some distal DHSs have particularly strong signals, similar to TSS DHS hits (**Fig. 1E**, **Fig. S5**). Significant DHSs were closer to TSSs than non-significant DHSs (**Fig. S5C-D**). While the majority of hits (68%) are promoters, which have the strongest effect sizes, the distal intergenic, intronic, exonic, and UTR hits have significant but weaker effect sizes in the same direction (**Fig. 1E, Fig. S5A-C**). For example, several DHS hits 10 kb - 1 Mb upstream of their putative target genes scored similarly to gRNAs that target the promoter of the same gene (**Fig. S6**). Some of the distal elements were previously validated in mice to control genes such as the oncogene *Lmo2* in erythroid cell lineages (Landry et al. 2009).

To identify epigenetic characteristics of DHSs that control cell fitness, we used ChromHMM genome annotations (Ernst and Kellis 2012) to identify classes of regulatory elements that were overrepresented in the enriched or depleted DHS hits (**Fig. 1F**). We observed hits in almost every class of annotation, including regions classified as polycomb-repressed.

Relative to all DHS sites, depleted DHS hits were overrepresented at active promoters and underrepresented at enhancers and CTCF sites. In contrast, enriched DHS hits were overrepresented at strong enhancers. An example of two enriched regions are intronic DHSs in the GMPR gene (**Fig. 1G**). GMPR encodes a protein responsible for purine nucleotide biosynthesis, and reduction of this gene has been linked to increased proliferation in some cell types (Wawrzyniak et al. 2013). The promoter and an upstream enhancer of GATA1 , as well as the HDAC6 promoter, are examples of significantly depleted regions and were previously identified as essential (**Fig. 1H**) (Fulco et al. 2016). While a variety of regions contribute to cell fitness, promoters and enhancers dominate the signal in this screen, as expected.

To better understand the characteristics that distinguish the significantly enriched or depleted gRNAs, we annotated each DHS with a selection of DHS-level features. Significant DHSs tended to be slightly longer, closer to TSSs, have nearest genes that were more highly expressed, have higher DNase accessibility, higher H3K27ac signal, and higher Hi-C contact frequency (**Fig. S7**). Additionally, genes nearest to significant DHSs were enriched for genes previously identified as essential (Hart et al. 2015; Wang et al. 2017) (**Fig. S7**). These features have been used previously to predict enhancer-gene interactions (Fulco et al. 2016, 2019), and support the power of this genome-wide screen to identify active regulatory elements associated with our cell fitness criteria.

For promoter hits, we compared our results to other studies of promoter inactivation (Horlbeck et al. 2016) or gene disruption (Lenoir, Lim, and Hart 2018; Wang et al. 2017) performed in K562 cells. The observed promoter hits are positively correlated (Pearson r = 0.64) with the promoter CRISPRi screen (Horlbeck et al. 2016) (**Fig. S8A, Table S2**). Overall, 250 genes were hits in all four studies, and our screen identified an additional 123 promoters as essential (**Fig. S8B**). Our genome-wide discovery screen identifies fewer promoters as essential, and consequently identifies fewer unique hits than previous genome-wide screens of coding genes, especially CRISPR-Cas9 knockout screens targeting exons (Wang et al. 2017; Lenoir, Lim, and Hart 2018).

### Validation of distal regulatory DHS hits by a secondary screen

We validated primary screen hits by performing a secondary cell fitness screen largely restricted to DHSs with significant gRNAs (**Fig. 1A**). Our screen was distinct from previous efforts in that most of our gRNAs targeted putative distal regulatory elements, not only gene promoters (Lenoir, Lim, and Hart 2018; Hart et al. 2014; Shalem et al. 2014; Horlbeck et al. 2016; Wang et al. 2017). To validate and characterize the effects of individual gRNA and DHS hits, we completed a validation screen of 232,346 unique gRNAs collectively targeting 8,845 unique DHSs (8,833 of which were also tested in the discovery screen) (**Fig. 1A, S8A-B**). All 8,833 DHSs from the discovery screen had at least one significant gRNA (FDR < 0.1) in the discovery screen, and 490 were significant in the stringent DHS-level analysis (FDR < 0.05, **Fig. S1, Table S2**). The DHSs included in the validation screen were largely distal, though some were annotated as promoters just downstream of the TSS (**Fig. S9A, B**).

Individual gRNA effects in the validation screen had a similar distribution as the discovery screen, both in terms of effect sizes and direction of effect on cell fitness (**Fig. 2A, Table S3**). The validation screen had more significant gRNA hits per DHS (**Fig. S9C**), suggesting that increasing the density of gRNAs tested per DHS from a mean of 10 to 26.2 improved detection of regulatory elements that impact cell fitness. This may be in part due to variation in the effect sizes of gRNAs targeting the same DHS (Thakore et al. 2015). However, the majority of DHSs still only had 1 individually significant gRNA (**Fig. 2B, Fig. S9C, Fig. S10A**). Unlike in the discovery screen, significant gRNAs in the validation screen were enriched for gRNAs with lower specificity, perhaps because the design prioritized additional gRNAs evenly spread across the entire DHS, rather than only the highest-specificity gRNAs (**Fig. S9D**).

**Figure 2:**
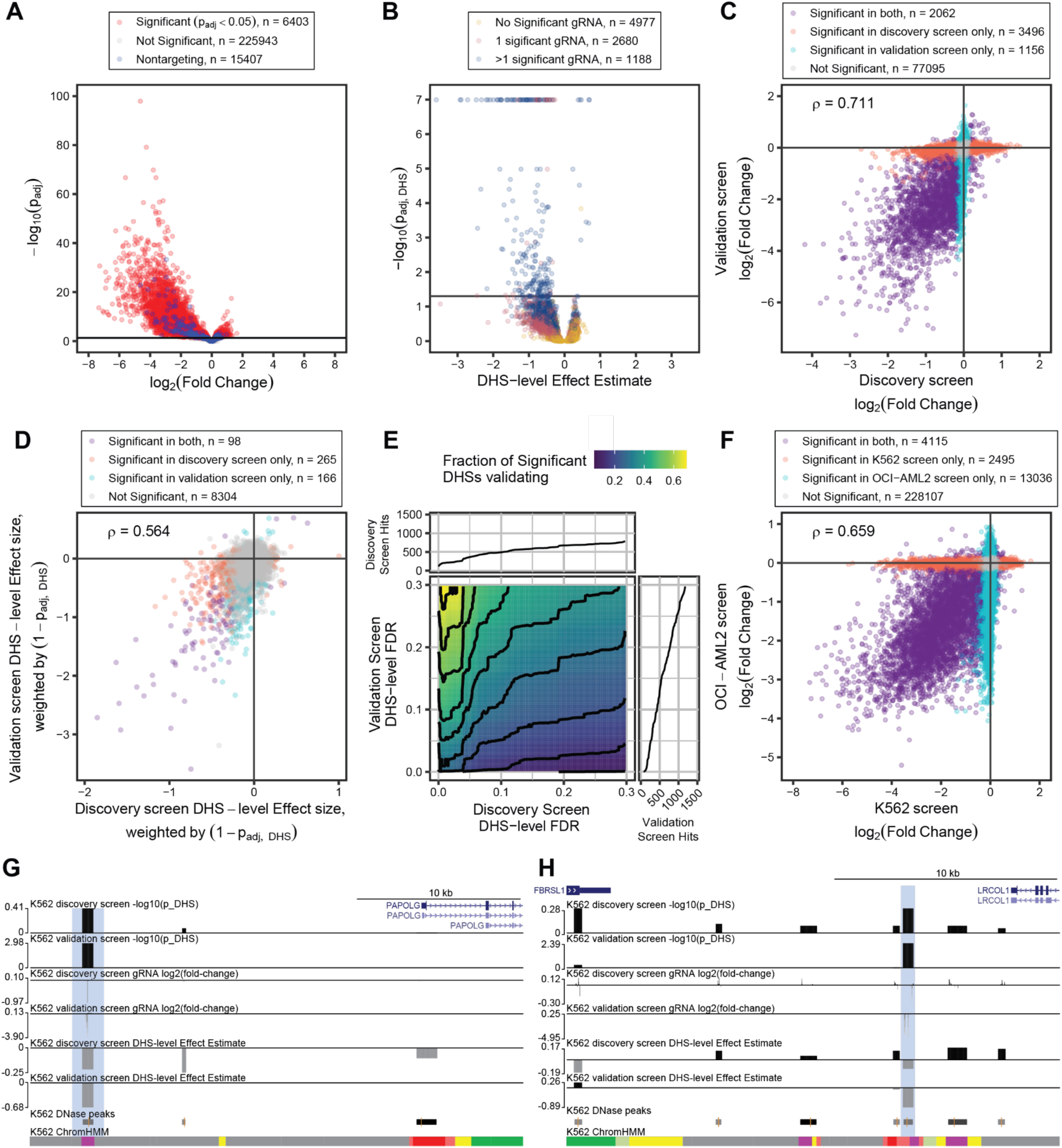
Validation of distal regulatory DHS hits using a secondary screen. **(A)** Significance of gRNA abundance changes relative to log_2_(fold-change) in the distal sub-library validation screen. **(B)** Significance of DHS effect on cell growth relative to aggregate effect size estimate, colored by the number of individually significant gRNAs in the DHS (FDR < 0.05). Horizontal line marks FDR = 0.05. **(C)** Correlation of gRNA-level effect sizes between the discovery and validation screens in K562s, colored by DHS-level significance in either screen (FDR < 0.1 in the initial screen, FDR < 0.05 in the validation screen). **(D)** Correlation DHS-level aggregate effect size estimates between the validation and discovery screens, weighted by DHS-level p-value. FDR cutoffs for significance are 0.05. **(E)** The fraction of DHSs found to be essential in the discovery screen that validate as essential in the secondary screen as a function of the false discovery rate (FDR) of both screens. The accumulation of elements called as significantly essential in either screen is shown in the marginal plots as empirical cumulative distribution function. Contour lines mark constant fractions. **(F)** Correlation of gRNA-level effect sizes in the secondary screen between OCI-AML2 and K562s. FDR cutoffs for significance are 0.05. **(G-H)** Representative browser tracks of DHS hits that displayed a denser tiling of gRNA hits in the distal validation screen near **(G)** *PAPOLG* and **(H)** *LRCOL1*. In both cases, more gRNAs with high abundance changes in these DHSs leads to a larger DHS effect estimate on growth and DHS significance.

After aggregation of gRNA signal to the DHS-level using nontargeting gRNAs as empirical negative controls, significant DHSs in the validation screen had similar distributions of significant scores and effect sizes on cell fitness as the discovery screen across promoters, intergenic regions, introns, exons, and UTRs (**Fig. 2B, Fig. S10**). Our analysis directly accounts for the number of gRNAs tested per DHS (**Fig. S9A**), so DHS significance remains FDR-controlled despite a variable number of tests per DHS.

To evaluate performance of single gRNAs, we characterized 83,809 individual gRNAs assayed in both the discovery screen and the validation screen. Effect sizes between the screens were highly correlated (Pearson ⍴ = 0.711, **Fig. 2C**), indicating a high level of replication between screens. This also supports that the gRNA effects in the initial genome-wide discovery screen are not specific to the clonal nature of K562-dCas9^KRAB^ cells used in it, since the validation screen used an independent polyclonal K562-dCas9^KRAB^ cell line. Restricting the analysis to gRNAs in DHSs that were significant in the discovery screen (**Fig. S11A**), either screen (**Fig. S11B**), or both screens (**Fig. S11C**) increases the correlation substantially, indicating the correlation is driven by the fitness effect of significant DHSs. The gRNA-level correlation remains very high over a wide range of DHS-level FDR thresholds, supporting reproducibility across screens (**Fig. S12A**).

Of the gRNAs tested in both screens, 5,558 were individually significant (FDR < 0.1) hits in the discovery screen and 2,172 reached the same level of significance in the validation screen (see marginal cumulative distribution functions in **Fig. S12B**). Of the 78,251 gRNAs that were negative in the discovery screen, 76,649 gRNAs were also negative in the validation screen (FDRs < 0.1). At a false discovery rate of 10% in both screens, the validation screen indicated gRNA-level sensitivity of 60% with 44% precision and 97% specificity in the initial discovery screen. The sensitivity, precision, and specificity vary as functions of the gRNA-level FDRs in both the validation and discovery screens (**Fig. S12B**). This high specificity is maintained across a range of gRNA-level false discovery rates. Sensitivity and precision have an expected inverse relationship across a range of FDRs. The modest sensitivity may indicate a substantial increase in power in the validation screen.

We also compared element-level performance between the two screens. The weighted DHS effect sizes correlated well across all DHSs tested (Pearson ⍴ = 0.564, **Fig. 2D**). Restricting to DHSs significant in the initial screen or both screens improved the correlation substantially (Pearson ⍴ = 0.63 and 0.72, respectively), indicating these are reproducible fitness-linked elements and validating both our screening and analysis methods (**Fig. S11D-F**). While the analysis statistically accounts for the number of gRNAs tested in each DHS, the power of the screen depends on the selected FDR (see the marginal cumulative distribution functions in **Fig. 2E, S11**). Similarly, the fraction of significant DHSs that are recovered in the secondary screen is dependent on the confidence-level threshold of both screens (**Fig. 2E**). Together, the secondary screen validates and expands the quantitative characterization of essential distal regulatory elements.

Of the 490 DHSs with significant (FDR < 0.1) effects in the discovery screen, 139 (28.3%) reached the same level of significance in the validation screen (see marginal cumulative distribution functions in **Fig. S12C**). Assuming the validation screen as ground truth, the validation indicates DHS-level sensitivity of 32% with 28% precision and 96% specificity (FDRs < 0.1) in the discovery screen, though these quantities vary as a function of the DHS-level FDRs (**Fig. S12C**). The low sensitivity of the discovery screen relative to the validation screen indicates a substantial increase in DHS-level power in the validation screen. Two examples of regulatory elements with more significant gRNAs with larger effect sizes, and a larger DHS effect size and significance in the validation screen, are intergenic regions annotated as inactive promoters upstream of *PAPOLG* (**Fig. 2G**) and *LRCOL1* (**Fig. 2H**).

### Identification of cell type-specific essential gene regulatory elements

To functionally assess the generalizability of essential regulatory elements across cell types, we re-purposed the validation gRNA library used on the chronic myeloid leukemia (CML) K562 cell line (**Fig. 2, S8**) to perform an additional screen in the acute myeloid leukemia (AML) cell line OCI-AML2 (**Table S4**). For the OCI-AML2 screen, inter-replicate gRNA abundance counts correlated more highly than in the K562 screen, with Pearson correlations above 0.91 among the dCas9^KRAB^ replicates, in comparison to correlations between 0.6 and 0.8 in the K562 screen (**Fig. S13**). This indicates a more homogeneous response to regulatory element perturbation in the OCI-AML2 cells.

Similar to the results for this library in K562 cells, we also detected depleted gRNAs with larger average effect sizes than enriched gRNAs (**Fig. S14A**), and significant gRNAs were enriched for low-specificity gRNAs (**Fig. S14D**). We detected 4,115 gRNAs significantly depleted in both cell types (**Fig. 2F**). We also detected 2,495 gRNAs significantly depleted only in K562 cells and 13,036 gRNAs significantly depleted only in OCI-AML2 cells (FDRs < 0.05), likely as a consequence of the tighter correlations between replicates in the OCI-AML2 screen (**Fig. S13**). These higher correlations and generally higher detection of significant gRNAs in the OCI-AML2 screen extended even to the nontargeting gRNAs; within the same pool of nontargeting gRNAs, 694 (0.3%) were significant in the OCI-AML2 screen, while only 207 (0.08%) were significant in the K562 screen (FDRs < 0.05). These nontargeting gRNAs estimate an empirical false discovery rate under 5%, with significant nontargeting gRNAs making up 3.1% and 4.0% of the pool of significant gRNAs in the K562 and OCI-AML2 screens, respectively (**Tables S3-4**). Across all the gRNAs in the library, effect sizes between cell types correlated slightly less well than between shared protospacers in the discovery and validation screens in K562 cells (**Fig. S15A-D**).

At the DHS-level, the OCI-AML2 screen also identified more DHSs that negatively affect cell fitness, finding 410 high-confidence fitness-linked regulatory elements (FDR < 0.05, **Fig. S14B-C**). The DHS weighted effect sizes correlated similarly between the cell types and within K562s (**Fig. S15E-H, S10D-F**). Within regions of strong overlap between cell types, there is consistency in the activity of shared gRNAs, with higher gRNA effect size correlations in regions with activity in both cell types (**Fig. S15**). While all the elements were chosen to have some activity in the original discovery screen in K562 cells, this data supports that 1) many of these elements have similarly strong fitness-linked activity across multiple cell types, and 2) some elements have stronger fitness-linked activity in certain cell types that indicates cell-type specific fitness pathways are being impacted. .

### Identification of regulatory element target genes using single cell gene expression

To empirically identify the target genes for the distal regulatory elements detected in these screens, we used Perturb-seq, a method that combines single cell RNA-seq readout with CRISPR screens (Adamson et al. 2016; Dixit et al. 2016; Datlinger et al. 2017; Xie et al. 2017; Gasperini et al. 2018). This allows the capture and quantification of all mRNA and gRNA identity on a per-cell basis, enabling the identification of genes that change in response to regulatory element perturbations. For this screen, polyclonal K562 cells constitutively expressing dCas9^KRAB^ were transduced with a library of 1,982 gRNAs targeting 350 DHSs (**Fig. S1, Table S5**). These gRNAs were the top 5 most significant per DHS from the validation screen, creating a compact and effective library from pre-screened gRNAs. Cells were collected and barcoded 7 days post-transduction to identify gene expression changes while minimizing library abundance changes due to fitness effects observed at later time points. The 350 DHSs targeted in this single-cell screen all negatively impacted cell fitness in the secondary validation screen and all had 1 or more significant gRNAs (**Fig. S16A**). Almost all of these DHSs were within 200kb of the nearest TSS, with only 2 DHSs > 1Mb away from a gene (**Fig. S16B**).

We transduced cells at a low MOI of ∼0.2 to ensure only 1 gRNA perturbation per cell. After selection and gRNA-cell assignment (**Methods**), gRNAs were represented by a median of 142 cells (**Fig. S17B**). differential gene expression was measured between cells with each gRNA and cells with nontargeting gRNAs (**Fig 3B**). Positive control gRNAs targeting the *HBG1/2* promoter formed highly significant gRNA-gene pairs downregulating *HBG1/2* and upregulating the other three globin genes (**Fig. S17C, Table S6-7**). Promoter control gRNAs targeting essential genes downregulated their target genes as expected (**Fig. S17C, Table S6-7**).

**Figure 3:**
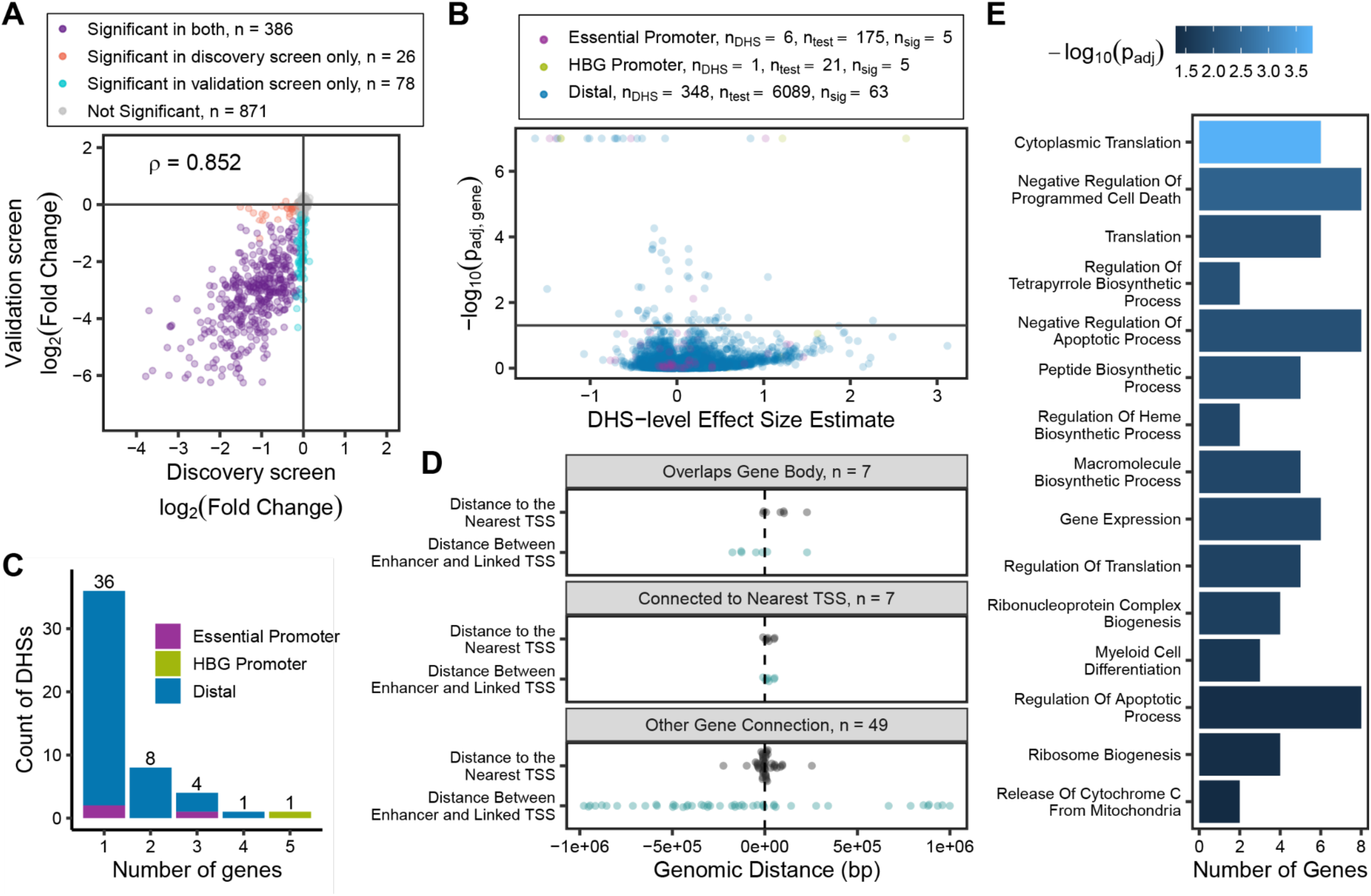
Identification of regulatory element target genes using single cell gene expression. **(A)** Correlation between the discovery and validation screen effect sizes for the 1,361 double-screened gRNAs of 1,733 gRNAs used in the one week screen. **(B)** Significance of DHS effect on gene expression relative to aggregate effect size estimate of the DHS-gene pair. Promoter controls for essentiality and expression (HBG) are colored separately. **(C)** Distribution of the number of genes linked to each DHS in the screen. The majority of DHSs were linked to only one gene, though 12 non-control DHSs were linked to multiple genes. The HBG promoter-targeting positive control led to changes in all globin genes regulated by the same locus control region. **(D)** Distribution of enhancer-promoter link length across 63 significant DHS-gene connections, divided by whether or not the enhancer is linked to either a gene it overlaps, the nearest TSS, or some other gene. **(E)** Gene ontology analysis on the 59 genes significantly linked to DHSs in this screen (FDR < 0.05), showing 15 significantly enriched pathways (FDR < 0.05).

Of the 6,089 tested DHS-gene pairs, we detect 63 significant regulatory interactions (FDR < 0.05, **Fig. 3B**). Since DHSs are tested for effects on any expressed gene (detected in >3 cells with a gRNA in the DHS, see Methods) in the ± 1 Mb window flanking that DHS, most pairs are not expected to be significant regulatory interactions. Collectively, we identified 59 genes that were affected by perturbing 46 unique regulatory elements. Approximately 60% of the regulatory elements (N = 34) affected a single gene, with the remainder impacting two or more genes (**Fig. 3C**). Generally, the experimentally identified enhancer-promoter links are much longer than the distance from a DHS to the nearest gene body or nearest transcription start site, supporting many of these are long range interactions (**Fig. 3D**). As expected, gene ontology analysis for the 59 target genes is significantly enriched for essential cell processes including translation, gene expression, and regulation of apoptosis (**Fig. 3E**).

This study identified 31 genes linked to distal enhancers in 33 DHS-gene connections that were not previously characterized as essential by three landmark studies (**Fig. S18A**) (Wang et al. 2017; Lenoir, Lim, and Hart 2018; Horlbeck et al. 2016). For twenty of these 33 DHS-gene connections that are unique to our study, dCas9^KRAB^-mediated repression of the DHS led to target gene upregulation, potentially explaining why promoter repression or gene knockout did not lead to a cell fitness phenotype in other studies (**Fig. S18B-C**). As expected, 25 of 28 distal links to previously identified essential genes downregulated the corresponding gene (**Fig. S18B-C**). The three genes that were downregulated but not known to be an essential gene are *FKBP8*, *RPL13A*, and *SESN2*. However, these are particularly long-range connections (100kb - 1Mb) and in two of those cases, the distal DHS also downregulated another previously identified essential gene. Thus these connections may be secondary or the result of an enhancer impacting multiple genes.

### Individual validations of select DHS-gene links

We next validated our screen results by testing a subset of individual gRNAs on the expression of their target genes using qRT-PCR. We tested 47 gRNAs across 10 DHSs affecting 13 genes at 1 week post-transduction of the gRNA (**Fig. 4A-C, Fig. S19, Table S8-10**). 39 out of the 40 (97.5%) significant individual gRNA-gene connections validated, and all 14 significant DHS-gene links validated. Some DHSs were significantly linked to multiple genes, such as chr6.4810 being linked to both MYB and AHI1 (**Fig. S19I, J**). In these cases, one link generally had a stronger effect size and higher significance in the screen, but both were validated by RT-qPCR. The fold changes in gene expression across significant DHS-gene links in the screen and fold changes in mRNA expression in the RT-qPCR correlated very strongly (Pearson r = 0.738, p < 10^!“#^), supporting the screening approach. For each DHS-gene pair, we also co-transduced the pooled collection of 4-5 gRNAs targeting that DHS. Each gRNA pool typically had similar effect sizes to that of the best individual gRNA **(Fig. 4B-C, Fig. S19**).

**Figure 4:**
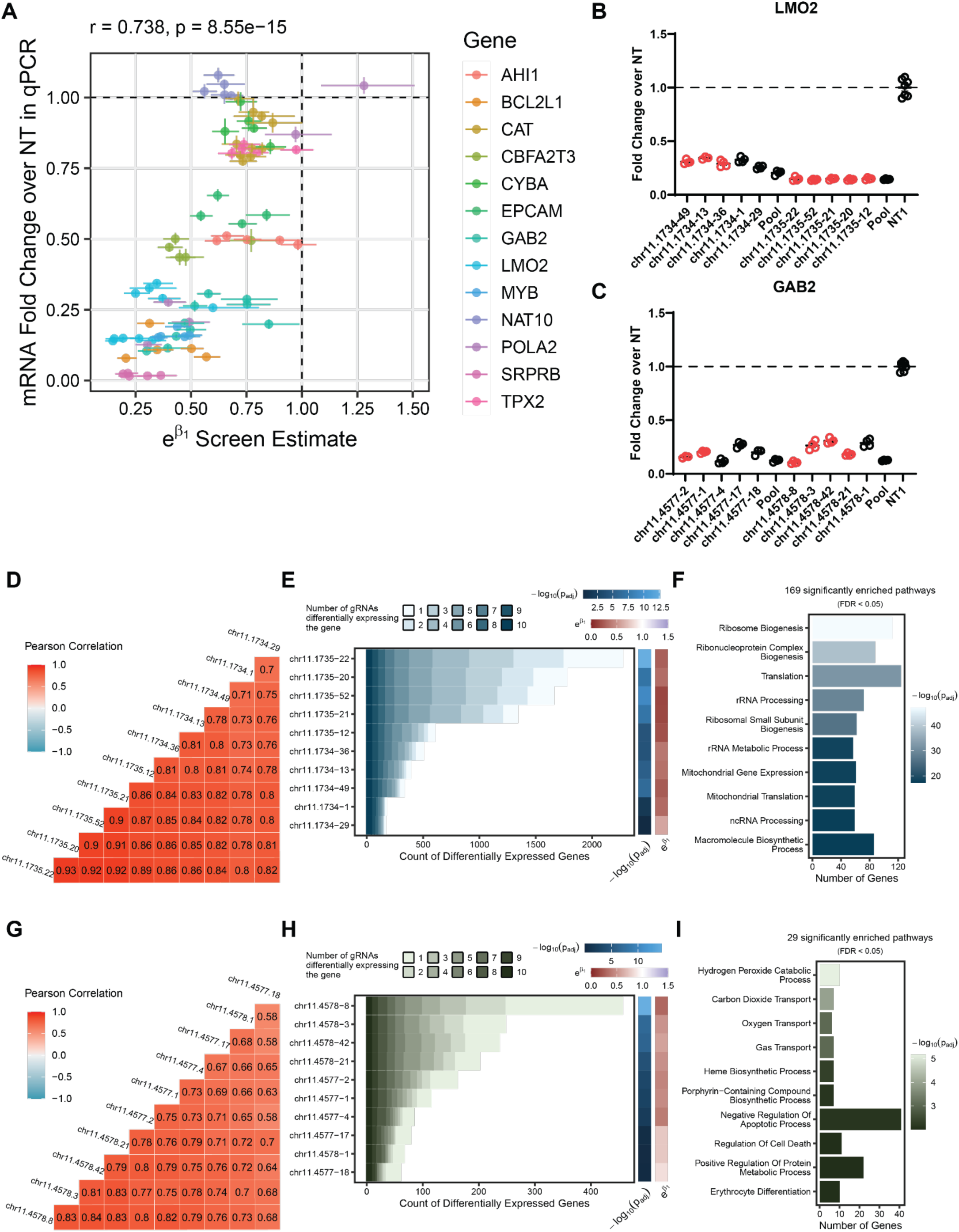
Single-Cell Transcriptional Validation. **(A)** Correlation of screen effect size and qPCR fold change for 47 validation gRNAs, colored by the gene of interest. Error bars in x indicate standard error estimates in the screen from the negative binomial model; error bars in y indicate standard error of 4 biological replicates from the qPCR. **(B-C)** qPCR validation of gRNAs changing **(B)** LMO2 and **(C)** GAB2 expression, respectively, displayed as mRNA fold change relative to NT control. Red points denote gRNAs with significant repression of the target gene in the single-cell screen. Black points denote non-significance in the single-cell screen link. gRNAs are ordered left-to-right in decreasing significance for this gene pairing in the single-cell screen. **(D-I)** transcriptome-wide correlation of differentially expressed genes (p<0.05) from the DHSs controlling **(D-F)** LMO2 and **(G-I)** GAB2 expression, respectively. **(D,G)** Transcriptome-wide correlation of the union set of differentially expressed genes across both DHSs **(E,H)** Counts of significantly differentially expressed genes for each gRNA, partitioned by the number of gRNAs detected as affecting the gene’s expression. Heatmaps on the right indicate magnitude of the gRNA’s effect on the target gene (𝑒^!!^ ) and the significance of that connection (𝑙𝑜𝑔_“#_ 𝑝_%&’_). **(F,I)** Gene ontology analysis across the union set of differentially expressed genes for the gRNAs in these DHSs.

The RT-qPCR assay generally achieved a higher degree of precision and confidence at the gRNA level (**Fig. S20A**) and across gRNAs within a DHS (**Fig. S20B**) than the scRNA-seq screen. Even in DHS-gene pairs with smaller effect sizes or only one or two individually significant gRNAs in the screen, the RT-qPCR effect sizes between gRNAs showed little variability (**Fig. 4B-C, Fig. S19E, H, J**).

### Transcriptome-wide effects of regulatory element perturbation

To measure transcriptome-wide effects of perturbing individual fitness DHS sites, we expanded our analysis of the single-cell screen data to include all genes. Perturbations of individual DHSs sometimes resulted in many differentially expressed genes associated with a variety of essential gene ontology terms, whereas others affected a much narrower set of genes (**Fig. 4D-I, Fig. S21**). Individual gRNAs corresponding to the same DHS showed very high correlations in global gene expression differences across their union set of differentially expressed genes (**Fig. 4D, G, Fig. S21**). Despite these individual gRNAs showing highly consistent effects on target gene expression in qPCR assays, the number of transcriptome-wide differentially expressed genes detected from each gRNA perturbation varied widely across a DHS, even in DHSs where all 5 gRNAs were significant (**Fig. 4E, H, Fig. S21**). For each DHS, the gRNA with the most differentially expressed genes often had the largest detected effect size on the primary target gene, and almost always had the most significant link to the target gene (see marginal heat maps, **Fig. 4E, H, Fig. S21A-L(ii)**). Occasionally, the majority of differentially expressed genes were driven by one gRNA, with poorer correlations between gRNAs that target the same DHS (**Fig. S21A, D, H, I**). The union sets of differentially expressed genes for each perturbation were associated with diverse pathways related to cell fitness, including some enriched for ribosome biogenesis, rRNA processing, and translation (**Fig. 4F, Fig. S21C-D, I, J, L**), while others were enriched for gas transport, heme synthesis, and myeloid differentiation (**Fig. 4I, Fig. S21E, F, G**). Some DHSs show fewer differentially expressed genes, identifying fewer or no significantly enriched gene ontologies (**Fig. S21 H, J, K**). These may represent perturbations with smaller downstream effects or fitness effects mediated through narrower pathways.

### Short-term CRISPRi screen reveals time-dependent effects of essential elements on cell fitness

In addition to the single cell gRNA screen, we used the same gRNA library to perform a very high coverage (∼3000X) one-week bulk gRNA abundance screen. At this high coverage, replicate correlations were very high (Pearson r > 0.98, **Fig. S22**), allowing precise detection of small abundance changes in the population over the one-week period. All of the gRNAs targeting essential gene promoter controls were significantly depleted (**Fig. S23B**), and all of the corresponding essential gene promoter elements were highly significant (**Fig. S23C**). Of the 350 distal DHSs screened, 341 had a significant effect on cell fitness at one-week post-transduction (FDRs < 0.05, **Fig. S23C**). Targeting the *HBG* promoter with the CRISPRi system increased cell fitness by a small but significant amount, presumably by eliminating the metabolic burden of producing a highly expressed gene (**Fig. S23A, C**). Across the 1,733 individual gRNAs screened in both the validation and 1-week screens, there was a strong and significant correlation (Spearman ⍴ = 0.714, p < 10^!“$^, **Fig. S23D**), showing the 1-week screen predicts screen results assessed at a later time point. The correlation at the element-level was more modest but still a significant association (Spearman ⍴ = 0.213, p = 7.7x 10^!#^, **Fig. S23E**). While the short-term screen was highly sensitive, fitness effect sizes of gRNAs (**Fig. S23D**) and elements (**Fig. S23E**) were more pronounced in the later time point of the validation screen. Many fitness-linked elements show strong and differentiating effects in the validation screen that they do not show after only one week.

### Functional validations of growth phenotype by competition assays

To further functionally characterize the top 58 gRNAs corresponding to 12 DHSs from the single-cell screen, we used a two-week cell growth competition assay to directly test whether silencing each distal regulatory element reduced cellular fitness (**Fig. 5A**). Cells expressing dCas9^KRAB^ were transduced with either a targeting or nontargeting gRNA and fluorescence marker and mixed 2 days post-transduction with untransduced cells. Fluorescent vs. non-fluorescent cell ratios were subsequently observed four times per day without antibiotic selection for 15 days (**Fig. 5A**). We first co-transduced the pooled collection of 4-5 gRNAs targeting each DHS, and show that all 12 pools reduced cellular fitness in comparison to a nontargeting gRNA (**Fig. 5B, Table S11**). We next tested individual gRNAs, and show that almost all reduced cellular fitness in comparison to a nontargeting gRNA. In some DHSs, variation in efficacy was observed for the different individual gRNAs throughout the time course. At the end of the time course (approximately matched to the time of the validation screen) this variation in the individual validations roughly corresponds to variation observed in the validation screen effect size (**Fig. S24, Table S11**).

**Figure 5:**
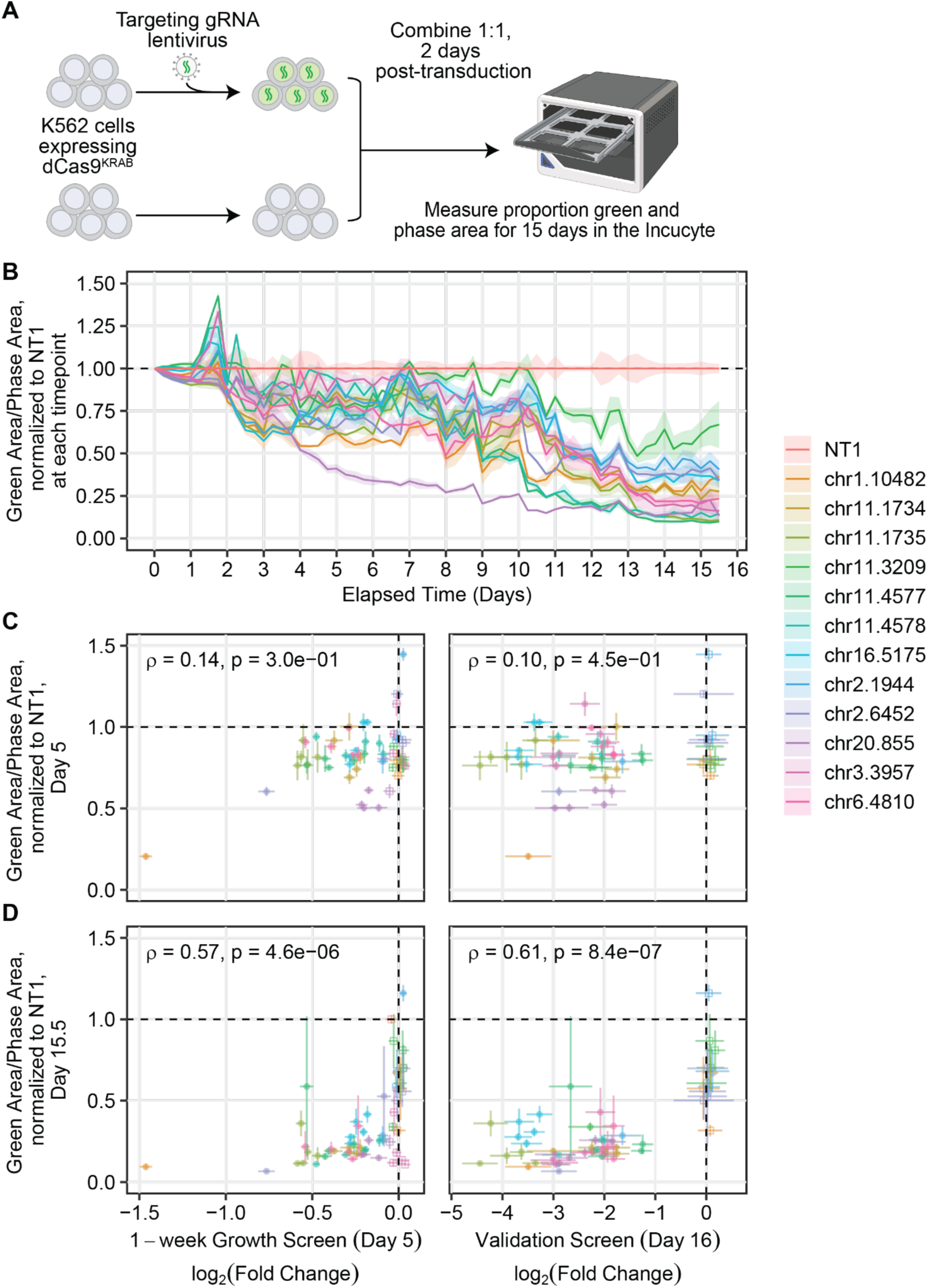
Competition Assay Validates Growth Phenotype. **(A)** Overview of competition assay method. Cells expressing dCas9^KRAB^ are transduced and combined 1:1 with untransduced cells at 10k/well 2 days post-transduction (day 0) to compete for 15 days in the Incucyte. **(B)** Pooled gRNA timecourses over the competition assay showing the decrease in green area relative to phase area across time, normalized to the nontargeting gRNA at each timepoint. **(C-D)** Comparison of cell fitness screen growth phenotypes (expressed as a log_2_(fold-change) in gRNA abundance) with time-matched individual validations at **(C)** day 5 in the competition assay or **(D)** the end of the competition assay. Error bars in x and y are standard error for the individual gRNA in the indicated screen or competition assay (n=4-5 technical replicates), respectively. Growth is expressed as a ratio of green area to phase area, relative to the initial timepoint for that sample and normalized to NT1 at each timepoint.

At 5 days in the competition assay, there are modest and insignificant correlations with the gRNA abundance changes in either the 1-week screen (Spearman ⍴ = 0.14) or the 16-day validation screen (Spearman ⍴ = 0.1 **Fig. 5C**). However, there is a much larger and statistically significant (p < 0.001) correlation between the growth disadvantage at the end of the competition assays and either the 1-week screen (Spearman ⍴ = 0.57) or the validation screen (Spearman ⍴ = 0.61, **Fig. 5D**). Notably, the high correlation between the phenotypes observed in the 1-week screen and at the end of the 2-week competition assay indicates that the high coverage 1-week bulk screen is reasonably able to predict the phenotype at a later time, though the relationship is not linear (**Fig. 5D**, left). In comparison, correlation of the competition assay at 16 days with the validation screen effect size reveals two distinct groups of gRNAs - those with weak and strong effect sizes (**Fig. 5D**, right). This indicates that the validation screen is well-powered to detect strong growth phenotypes, but is unable to detect more subtle changes in cellular fitness.

Together, the competition assay time course experiments support that some gRNAs corresponding to an individual fitness-related DHS have a stronger effect than others and that the growth screens quantitatively measure reproducible phenotypic variation between gRNAs within the same DHS.

## Discussion

Human phenotypes and disease susceptibilities are linked to the function of gene regulatory elements and associated genetic variation in the non-coding genome (Tam et al. 2019; Maurano et al. 2012). There is strong rationale for cancer susceptibility and progression to be similarly dictated by the non-coding genome, and a few discrete examples of this mechanism have emerged (Khurana et al. 2016; Laumont et al. 2018; Orlando et al. 2018; Hayward et al. 2017). However, cancer genetics and the discovery of oncogenic driver mutations has historically been limited to analysis of protein coding sequences because (i) whole-genome sequencing of primary tumors is costly, and (ii) our functional understanding of noncoding genetic variation is still in its infancy. The first limitation is being addressed by lower costs of whole genome sequencing and growing databases of thousands of whole genomes from tumor and healthy control tissue (Rheinbay et al. 2020; Corces et al. 2018). However, these advances also emphasize the urgency in addressing the second limitation, which has recently become a tractable challenge due to the advent of CRISPR-based screens of non-coding regulatory elements in their endogenous chromosomal context (Wang et al. 2014; Shalem et al. 2014; Gilbert et al. 2014; Thakore et al. 2015; Canver et al. 2015; Fulco et al. 2016; Sanjana et al. 2016; Klann et al. 2017). Knowing the location of regulatory elements that affect cell growth is a critical first step to find additional noncoding cancer driver mutations (Lawrence et al. 2014; Rheinbay et al. 2020).

This study is a significant step towards addressing these limitations and realizing the potential of whole genome sequencing for cancer biology. We describe a systematic genome-wide screen of all putative regulatory elements in a commonly used cancer cell line and quantify the impact of these elements on cell fitness. We identified >1,000 regulatory elements that have negative or positive impacts on cellular viability and/or proliferation, and characterizing a subset of these, reporting 63 distal element-gene links that contribute to this phenotype. These data provide a rich resource of regulatory element function and connection to target genes that will be broadly useful for understanding gene network regulation and the mechanisms of non-coding element control on gene expression. We expect these characterizations that relate the non-coding genome to cell fitness will identify functional noncoding sequence variants that contribute to cell growth phenotypes, including oncogenesis, as well as erythrocyte differentiation and cell metabolism. These functional annotations also complement the growing body of chromatin conformation maps that provide structural relationships between regulatory elements and genes (Kempfer and Pombo 2020).

Importantly, our work provides evidence that while some fitness-linked regulatory elements have similar effects across multiple cell types, that some elements have more potent effects on fitness in certain cell lineages. This study provides a blueprint for executing similar studies in other healthy and cancer cell types, genetic backgrounds, environmental conditions, or pharmacologic treatments. In the future, we expect that this approach will identify element-gene relationships and fitness pathways that are common or unique to certain cancer cell types or conditions. These will provide important information on new therapeutic targets that target unique fitness pathways in certain cancer cell types.

A challenge to implementing genome-wide screens of the non-coding genome is the sheer scale of the experiment, which is dictated by the number of putative elements in any cell type and the required numbers of gRNAs per element and cells per gRNA. As the field of CRISPR-based screens grows, more efficient and sensitive screening methods will be needed. For example, relatively little is known about which key gRNA attributes contribute to effective perturbation of distal regulatory elements. We expect that the knowledge gained from thousands of gRNAs that impact cellular growth from distal regulatory elements will facilitate the design of more compact and robust libraries, and enable similar genome-wide screens in cell lines or primary cells that are more difficult to culture at scale.

Many epigenetic modifying drugs used as potential cancer treatments cause widespread changes throughout the genome (Frank et al. 2016; Szyf 2009). However, it is currently unclear what subset of gene regulatory elements drive drug response. Using maps of essential regulatory elements in conjunction with the epigenetic profiles of cells after drug treatment could help identify modifications to specific gene regulatory elements necessary and sufficient for drug response. This may ultimately inform the development of safer and more specific cancer therapies.

Interestingly, one of the loci with the strongest effect on cellular proliferation was the *LMO2* locus. This locus is also the location of retroviral insertions in gene therapy patients which lead to increased expression of LMO2 via viral enhancer elements and ultimately to leukemia (Hacein-Bey-Abina et al. 2003). Better understanding the regulatory landscape of these and other types of regions will help elucidate mechanisms of aberrant gene expression and tumorigenesis that will ultimately also inform design, safety monitoring, and regulation of emerging classes of genetic medicines such as gene therapy and genome editing. Therefore we anticipate this approach will be a valuable resource to diverse fields of the biomedical research community.

## Materials and Methods

### Plasmids

The lentiviral dCas9-KRAB plasmid (Addgene #83890) was generated by cloning in a P2A-HygroR (APH) cassette after dCas9-KRAB using Gibson assembly (NEB, E2611L). The lentiviral gRNA expression plasmid was cloned by combining a U6-gRNA cassette containing the gRNA-(F+E)-combined scaffold sequence (B. Chen et al. 2013) with an EGFP-P2A-PAC cassette into a lentiviral expression backbone (Addgene #83925) using Gibson assembly. Individual gRNAs were ordered as paired oligonucleotides (IDT-DNA), phosphorylated, hybridized, and ligated into the EGFP gRNA plasmid using BsmBI sites.

### Cell Culture

K562 and HEK293T (for lentiviral packaging) cells were obtained from the American Tissue Collection Center (ATCC) via the Duke University Cancer Center Facilities. OCI-AML2 cells were gifted from Anthony Letai at Dana Farber Cancer Institute. K562 and OCIAML2 cells were maintained in RPMI 1640 media supplemented with 10% FBS and 1% penicillin-streptomycin. HEK293T cells were maintained in DMEM High Glucose supplemented with 10% FBS and 1% penicillin-streptomycin. All cell lines were cultured at 37 °C and 5% CO2.

For the genome-wide discovery screen, a clonal K562-dCas9^KRAB^ cell line was used, and was generated by transduction of dCas9-KRAB-P2A-HygroR lentivirus with polybrene at a concentration of 8 µg/mL. Cells were selected 2 days post-transduction with Hygromycin B (600 µg/mL, ThermoFisher, 10687010) for 10 days followed by sorting single-cells into 96-well plates with a SH800 sorter (Sony Biotechnology). Individual clones were grown and stained for dCas9^KRAB^ with a Cas9 antibody (Mouse mAb IgG1 clone 7A9-3A3 Alexa Fluor 647 Conjugate, Cell Signaling Technologies, 48796) to assess protein expression. Briefly, 1×10^6^ cells were harvested and washed once with 1X FACS buffer (1% BSA in PBS). The cells were then fixed and permeabilized for 30 minutes at room temperature with 500 µL of fixation and permeabilization buffer (eBioscience Foxp3/TF/nuclear staining kit, ThermoFisher, 00-5523-00). Next, 1 mL of permeabilization buffer was added and cells were pelleted (600 RCF for 5 min) and washed again in 1 mL of permeabilization buffer. Cells were pelleted again and resuspended in 50 µL of permeabilization buffer with 2% mouse serum (Millipore Sigma, M5905) to block for 10 minutes at room temperature. Following blocking, 50 µL of permeabilization buffer with 2% mouse serum and 1 µL of Cas9 antibody was added and allowed to incubate for 30 minutes at room temperature. Following incubation, 1 mL of permeabilization buffer was added, cells were pelleted and washed once more with 1 mL of permeabilization buffer. Finally, cells were resuspended in 1X FACS buffer for analysis. Each clone was analyzed using an Accuri C6 flow cytometer (BD Biosciences). A clone was selected based on high and uniform expression of dCas9^KRAB^ and expanded for further use.

For the secondary sub-library screens, we used polyclonal K562 and OCI-AML2 cell lines that express the dCas9^KRAB^ repressor. Polyclonal lines were used to account for possible hits in the first screen that could be specific to the clonal line used. K562 and OCI-AML2 cells were transduced with dCas9-KRAB-P2A-HygroR lentivirus with polybrene at a concentration of 8 µg/mL. At two days post-transduction, cells were selected for 10 d in Hygromycin B (600 µg/mL). Following selection, polyclonal cells were stained to detect expression of dCas9^KRAB^ protein as described above.

### gRNA Library Design

DNase I hypersensitive sites (DHSs) for the K562 cell line were downloaded from www.encodeproject.org (ENCFF001UWQ) and used to extract genomic sequences as input for gRNA identification. We used the gt-scan algorithm to identify gRNA protospacers within each DHS region and identify possible alignments to other regions of the genome (O’Brien and Bailey 2014). The result is a database containing all possible gRNAs targeting all targetable DHSs in K562 cells and each gRNA’s possible off-target locations. gRNAs were selected based on minimizing the number of off-target alignments. For the initial genome-wide library, 1,092,706 gRNAs were selected, targeting 111,756 DHSs (269 DHSs contained no NGG SpCas9 PAM), limited to a maximum of 10 gRNAs per DHS (mean, 9.77 gRNAs per DHS, **Fig. S1B**). Excluding exactly duplicated protospacer sequences, the results are presented as 1,084,704 gRNAs targeting 111,619 DHSs (**Table S1**).

For the second sub-library 234,593 gRNAs were selected, targeting 8,850 distal DHSs. Excluding exactly duplicated sequences, the results are presented as 232,346 unique gRNAs collectively targeting 8,845 unique DHSs (**Table S3**). All 8,833 DHSs with results in the discovery screen had at least one significant gRNA (FDR < 0.1), and there were 12 DHSs dropped out of the initial screen included in the validation. The sublibrary targeted mostly distal non-promoter hits (>3kb from TSS) identified in the first screen, though some sequences just downstream of TSSs annotated as promoters were included (**Fig. S9A-B**).

For each DHS, gRNAs were chosen to be spread evenly across the region by dividing each DHS into bins of 100 bp and selecting up to 7 gRNAs per bin. The gRNAs for each bin were selected in order by the fewest number of off-target alignments calculated by gt-scan. 15,407 non-targeting gRNAs were designed as previously described (Horlbeck et al. 2016). A larger number of gRNAs per DHS were designed in the second screen (mean of 26 per DHS, **Fig. S9A**) compared to the first screen (maximum of 10 per DHS).

For the single-cell screen, 350 distal (>3kb from TSS) DHSs were chosen from the distal sublibrary screen. To exclude DHSs with dual effects, DHSs were required to have a consistent direction of effect between the initial screen and the sublibrary screen without significant gRNAs with opposing effects. The top 350 DHSs meeting these criteria were chosen by DHS-level effect size; the top 5 most significant gRNAs from the distal sublibrary screen were chosen to be synthesized in a smaller pool for a total of 1,733 targeting gRNAs. 1,361 of these targeting gRNAs were included in the initial screen, and their effects between the two screens correlated well, Pearson’s ⍴ = 0.852 (**Fig. 3A**). Not all gRNAs chosen for this screen were significant in the distal sublibrary screen, but all DHSs had negative effects. The library also contained 30 gRNAs targeting 6 essential genes as positive growth controls, 20 HBG promoter-targeting gRNAs as positive controls, and 199 (10%) non-targeting gRNAs as negative controls (**Table S5**).

All libraries were synthesized by Twist Biosciences and the oligo pools were cloned into the lentiviral gRNA expression plasmid using Gibson assembly as previously described (Klann, Black, and Gersbach 2018). Briefly, oligo pools were amplified across 16 PCRs (100 ng oligo per PCR) for 10 cycles using Q5 2X master mix and the following primers:

Fwd: 5’-TAACTTGAAAGTATTTCGATTTCTTGGCTTTATATATCTTGTGGAAAGGACGAAACACCG Rev: 5’-GTTGATAACGGACTAGCCTTATTTAAACTTGCTATGCTGTTTCCAGCATAGCTCTTAAAC

Pools were gel purified (Qiagen, 28704) and used to assemble plasmid pools with Gibson assembly (NEB, E2611L). Pools were assembled across 16 Gibson assembly reactions (∼900 ng backbone, 1:3 backbone to insert) for the first screen, 4 reactions for the second sub-library screen. For the single-cell library, only 10 ng of the oligo pool was amplified for 10 cycles using Q5 2X master mix before gel purification and Gibson assembly (∼840 ng backbone, 1:3 backbone to insert) in one reaction throughout.

### Lentivirus Production

The lentivirus encoding gRNA libraries or dCas9^KRAB^ was produced by transfecting 5×10^6^ HEK293T cells with the lentiviral gRNA expression plasmid pool or dCas9^KRAB^ plasmid (20 µg), psPAX2 (Addgene, 12260, 15 µg), and pMD2.G (Addgene, 12259, 6 µg) using calcium phosphate precipitation (Salmon and Trono 2006). After 14-20 hours, the transfection media was exchanged with fresh media. Media containing lentivirus was collected 24 and 48 hours later. Lentiviral supernatant was filtered with a 0.45 µm CA filter (Corning, 430627). The dCas9^KRAB^ lentivirus was concentrated 20X the initial media volume using Lenti-X concentrator (Clontech, 631232), following manufacturer’s instructions. The lentivirus encoding gRNA libraries was used unconcentrated.

The titer of the lentivirus containing either the genome-wide library or distal sub-library of gRNAs was determined by transducing 5×10^5^ cells with varying dilutions of lentivirus and measuring the percentage of GFP-positive cells 4 days later using the Accuri C6 flow cytometer (BD Biosciences).

To produce lentivirus for the scRNA-seq screen, 10×10^6^ HEK293T cells were transfected with 4300 ng of the gRNA plasmid pool, 3250 ng PMD2.G and 9750 ng psPAX2 with lipofectamine 3000. The titer was determined by transducing ∼400k cells at the same seeding density as the screen with varying dilutions of lentivirus and measuring the percentage of GFP-positive cells 2 days later using the BD Accuri C6 flow cytometer.

To produce lentivirus for individual gRNA validations, 1.2×10^6^ HEK293T cells were transfected with gRNA plasmid (500 ng), psPAX2 (1.5 µg), and pMD2.G (500 ng) using 7.5 uL Lipofectamine 3000 and 5 uL P3000 reagent. Cell culture for virus production was all done in Opti-MEM supplemented with 5% FBS, 1X GlutaMAX (Gibco 35050061), 1mM sodium pyruvate and 1x MEM Non-essential amino acids (Gibco 11140050). After 4-6 hours, transfection media was exchanged with fresh media. Lentiviral supernatant was harvested 24 and 48 hours later, filtered through a 0.45 µm CA filter, and concentrated using Lent-X concentrator to 20X the initial media volume.

### Lentiviral gRNA Screens

For the first genome-wide screen, 1.7×10^9^ cells were transduced with the gRNA library during seeding in 3 L spinner flasks across 4 replicates for controls (K562 cells without dCas9^KRAB^) and 4 replicates for dCas9^KRAB^-expressing cells. For sub-library screens, 4.17×10^8^ cells were transduced during seeding in 500 mL spinner flasks across 4 replicates for both controls and dCas9^KRAB^-expressing cells. Cells were transduced at a multiplicity of infection (MOI) of 0.4 to generate a cell population with >80% of cells harboring only 1 gRNA and 500-fold coverage of each gRNA library. After 2 days, cells were treated with puromycin (Millipore Sigma, P8833) at a concentration of 2 µg/mL. Cells (control and dCas9^KRAB^-expressing) were selected for 7 days and allowed to grow for a total of 16 days (including 7 days of selection, or ∼14 doublings). Cells were passaged to ensure at least 500X fold coverage of the gRNA library to maintain representation. After culturing, for the genome-wide screen, 5.5×10^8^ K562 cells were harvested for genomic DNA isolation. For the sub-library distal screens in K562 cells or OCI-AML2 cells, 1.5×10^8^ cells were harvested. Genomic DNA was harvested from cells as described by Chen and Sanjana et al.(S. Chen et al. 2015).

For the one-week cell fitness screens, 32.8×10^6^ polyclonal K562s constitutively expressing dCas9^KRAB^ were transduced without polybrene at an MOI of 0.2 across 8 replicates at 3000X coverage for both controls and dCas9^KRAB^-expressing cells. Cells were maintained in 2 ug/mL puromycin (Thermofisher A1113803) from 48 hours post-transduction until collection at 7 days post-transduction (5 additional days of growth). Genomic DNA was harvested from ∼6.8×10^6^ cells per replicate by overnight incubation at 55 C in 2 mL of NK lysis buffer (50 mM Tris, 50 mM EDTA and 1% SDS) with 10 uL of 20 mg/mL Proteinase K (NEB P8107S). The next day, samples were shaken at 37 C after the addition of 10 uL of 10 mg/mL RNAse A (Qiagen 19101) for 3-4 hours. Samples were cooled on ice before adding 0.67 mL of chilled 7.5M ammonium acetate, vortexed at high speed for 30s and centrifuged at >4000g for 10 min. The supernatant was decanted into new tubes and gDNA was precipitated out with 3 mL of 100% isopropanol. The tubes were inverted 50 times to rinse and centrifuged at >4000g for 10 min. The supernatant was discarded; tubes were inverted 10 times after addition of 3 mL of freshly prepared 70% ethanol to rinse the pellet, then centrifuged at >4000g for 1 min to re-pellet. The supernatant was removed; the gDNA pellet was allowed to air dry for 30 minutes, then resuspended in 200 uL of water. All gDNA samples were quantified after purification using the Qubit dsDNA Broad Range Assay Kit (ThermoFisher, Q32850).

### Single-Cell RNA-seq Screen

At collection of the 1-week screen, cells were barcoded with the 10X 5’ HT v2 chemistry for single-cell RNA-sequencing. Fifteen lanes were collected from the first replicate of the low MOI bulk screen for a low MOI single-cell RNA-seq screen. Gene expression and gRNA libraries were generated and sequenced on a NovaSeq S4 flow cell according to the corresponding 10X 5’ HT v2 protocol.

### Genomic DNA Sequencing

To amplify the genome-wide gRNA libraries from each sample, 5.25 mg of genomic DNA (gDNA) was used as template across 525 x 100 µL PCR reactions using Q5 2X Master Mix (NEB, M0492L). For the distal sub-library screens, 1.2 mg of gDNA was used as template across 120 PCR reactions using Q5 2X Master Mix. Amplification was carried out following the manufacturer’s instructions using 25 cycles at an annealing temperature of 60 °C using the following primers:

Fwd 5’-AATGATACGGCGACCACCGAGATCTACACAATTTCTTGGGTAGTTTGCAGTT

Rev 5’-CAAGCAGAAGACGGCATACGAGAT (6 bp index sequence) GACTCGGTGCCACTTTTTCAA

Amplified libraries were purified using Agencourt AMPure XP beads (Beckman Coulter, A63881) using double size selection of 0.65X and then to 1X the original volume. Each sample was quantified after purification using the Qubit dsDNA High Sensitivity Assay Kit (ThermoFisher, Q32854). Samples were pooled and sequenced on a HiSeq 4000 or NovaSeq 6000 (Illumina) at the Duke GCB sequencing core, with 21 bp single read sequencing using the following custom read and index primers:

Read1 5’-GATTTCTTGGCTTTATATATCTTGTGGAAAGGACGAAACACCG Index 5’-GCTAGTCCGTTATCAACTTGAAAAAGTGGCACCGAGTC

For the single-week bulk screen, the same primers were used to amplify ∼40 ug of gDNA for 18 cycles over 24 reactions (1.75 ug per 100 uL reaction) using Q5 2X Master Mix. Amplified libraries were purified using double size selection of 0.6X and 1.8X the original volume using AMPure XP beads, and quantified as above. Sequencing was carried out using a NextSeq 550 (Illumina) at the Duke sequencing core with the custom primers listed above.

### Data Processing and Differential Expression Analysis of Bulk gRNA libraries

To identify and quantify the effects of regulatory element perturbation on cell fitness, we compared gRNA abundance before and after cell growth. FASTQ files were aligned to custom indexes (generated from the bowtie2-build function) using Bowtie2 (Langmead and Salzberg 2012) (options `--norc -p 24 --no-unal --end-to-end --trim3 6 -D 20 -R 3 -N 0 -L 20 -a` for the initial screen and sublibrary, `--norc -p 8 --no-unal -- end-to-end --trim3 2 -D 16 -R 3 -N 1 -L 20` for the single week bulk screen). Counts for each gRNA were extracted and used for further analysis. The third replicate of the genome-wide discovery screen was excluded from downstream analysis because its counts correlated poorly with the other 3 replicates. All gRNA enrichment analysis was performed using

R. For differential expression analysis, the DESeq2 package was used to compare between dCas9^KRAB^ and control (no dCas9^KRAB^) conditions for each screen. Log2 fold-change values were shrunk towards zero using the adaptive shrinkage estimator from the ‘ashr’ R package (Stephens 2017). Both the initial DHS data and gRNA libraries contained exact sequence duplicates; 8,002 gRNA IDs were sequence duplicates and removed from downstream analysis.

We used a novel robust rank aggregation technique to generate an FDR-controlled DHS-level significance score. First, DESeq2 screen p-values were transformed against a null empirical cumulative distribution function (ECDF) generated using the non-targeting or negative control gRNAs. This ensures that the null p-values within the experiment are uniformly distributed. As library size constraints prevented the use of non-targeting gRNAs in the initial discovery screen, negative control DHSs were chosen from the library. Negative control DHSs were chosen as the closest DHS less than 1 kb in length within 3 kb of any given non-essential gene TSS. From a list of 927 non-essential genes (Hart et al. 2014), this amounted to 2,878 gRNAs within 292 DHSs (**Fig. S3A-B**). By construction, these non-essential gene promoter DHSs are insignificant (**Fig. 1D, Fig. S3B**).

Transformed p-values were ordered within each DHS. Under the null hypothesis, p-values are uniformly distributed such that their order statistics follow a beta distribution with parameters dependent on the rank within the DHS and the number of gRNAs tested. P-values from those beta-distributions were compared to p-values from 10 million null simulations to construct DHS-level p-values for each gRNA. The minimum of the top 30% of the order statistics was taken to be the DHS-level p-value. In the initial genome-wide discovery screen, this means that significant DHSs have “surprisingly” small p-values in at least one of the first three order statistics by construction. There is no significant difference between average gRNA p-values across all orders between the negative controls and the full set of DHSs tested, likely because most DHSs do not affect cell fitness (**Fig. S25A**). Interestingly, significant DHSs have, on average, smaller p-values than negative control or insignificant DHSs all the way up to the 10th order statistic, despite only having one significant gRNA in most cases (**Fig. S25B**, Wilcox test with Bonferroni correction, p<2e-16 across all orders). There is no significant difference between the negative controls and non-significant DHSs, on average, across all orders (**Fig. S25B**).

In the 1-week screen, nontargeting gRNAs appeared enriched because so many of the targeting gRNAs had an effect on cell fitness and were depleted. To correct for this, all the gRNA abundance log_2_(fold-change) values were shifted by the nontargeting gRNA mean log_2_(fold-change), and significances for each gRNA were re-calculated on based on the shifted log_2_(fold-change). Shifted values are presented in **Fig. S23**.

To summarize enrichment or depletion across a DHS in the discovery and validation screens, we generated a weighted average of the log_2_(fold-change) in gRNA abundance across the gRNAs within a DHS. Two weighted averages were generated per DHS: the first weighting the log_2_(fold-change) by their lower tail transformed p-values, giving greater weight to large positive fold changes, and the second weighting by the upper tail transformed p-values, giving greater weight to large negative fold changes. The absolute maximum of the two was chosen as the DHS-level effect estimate. Notably, this use of the absolute maximum rather than the total average means small gRNA-level deviations from 0 will push the DHS-level effect size estimate further from 0, rather than closer to 0.

### Data Processing and Differential Expression Analysis of Single-Cell RNA-seq Screen

Sequencing data from transcriptome and gRNA libraries were processed using 10X Genomics CellRanger 7.1.0. Reads were first demultiplexed using the `mkfastq` command from 10X Genomics Cell Ranger 7.1.0 with the default configuration; transcript and gRNA (feature) counts were generated using the count command and the CellRanger GRCh38-2020-A hg38 reference dataset. At this point, 14 of the 15 lanes in the experiment were aggregated together using CellRanger `aggr`; one lane was pre-processed separately because it suffered from very low CRISPR library counts (median of 70 UMIs per cell). The bulk of the data consisted of 265,949 cells with a per cell median of 10,555 transcriptomic UMIs across 3,666 genes, and a median of 1,852 gRNA UMIs per cell. The single lane processed separately consisted of 21,115 cells with a median of 12,011 transcriptomic UMIs across 3,808 genes per cell.

Using Seurat v4.3.0, cells with >20% of mitochondrial UMI counts, <1,000 transcript UMIs, or < 500 genes were filtered out. Cells with transcript UMIs more than 2 IQRs beyond the 75% percentile of the dataset were also filtered out. At this point, transcriptomic UMI counts were normalized using `sctransform` with 2,000 genes and 10,000 maximum cells (Hafemeister and Satija 2019).

From the filtered data, we assigned gRNAs to cells using CLEANSER (Liu et al. 2025). CLEANSER is a mixture model that estimates posterior probabilities of the presence of each gRNA in each cell from the gRNA UMI counts in each cell, accounting for the distribution of the particular gRNA across cells and the particular cell. gRNA-cell pairs with probabilities > 0.9 were assigned and cells without gRNAs at that cutoff were removed. CLEANSER was run separately on the lane with very few CRISPR library reads, successfully assigning gRNAs to 16,174 (79%) of cells, despite having a median of only 70 gRNA UMIs per cell. After assignment, this lane was merged with the other data. This left 239,978 high quality cells. Cells with no gRNA assigned were discarded. The gRNA-containing cells had an average of 1.33 gRNAs per cell, confirming the low MOI (**Fig. S17A**). Each gRNA was represented by a median of 142 cells (**Fig. S17B**), with a total of 239,978 cells with gRNAs assigned.

To increase statistical power to detect changes in gene expression, differential expression tests for each gRNA were limited to genes in a 2 megabase window centered on the gRNA midpoint. Genes were identified using the Ensembl v104 reference and tested for differential expression using the `MASS:glm.nb` negative binomial model. The set of cells with each gRNA was compared to the set of cells with only non-targeting gRNAs to fit a coefficient describing the effect of the gRNA on gene expression (𝛽_“_) and a coefficient describing the basal gene expression (𝛽_%_) (**Table S6**). gRNA-gene pairs expressed in less than 3 cells were not excluded from the analysis and not tested. The union set of all genes within the testing windows of all gRNAs was used to run the same analysis for non-targeting guides. gRNA-gene pairs were detected in a median of 34 cells (**Fig. S17D**) and DHS-gene pairs were detected in a median of 161 cells (**Fig. S17E**), with a median coverage of 2,916 cells per DHS (**Fig. S17F**). gRNA-gene pair results were aggregated to DHS-gene pair results using the same aggregation approach as above. The non-targeting gRNA-gene pair results were used to form a midpoint-interpolated empirical cumulative distribution function which was in turn used to adjust the p-values of the test gRNA-gene pairs before aggregation. Effect sizes to summarize gene activation or repression were generated by applying the same weighted average technique as above to the 𝛽_“_coefficients from the negative binomial model.

### Individual gRNA Validations using qRT-PCR and Growth Competition Assays

Validation of individual gRNAs in distal (non-promoter) putative regulatory elements focused on the 12 DHSs forming the top 19 DHS-gene connections from the single-cell screen. All 5 protospacers from the single-cell screen were ordered as oligonucleotides from IDT and cloned into a lentiviral gRNA expression vector as described earlier. The same modified cell lines used in the corresponding screen were used for the individual gRNA validations. The cells were transduced with individual gRNAs and after 2 days were selected with puromycin (2 µg/mL) for 7 days.

For all screen validations by qRT-PCR, mRNA expression analysis was done in biological quadruplicate. Total mRNA was harvested from cells using the MagMAX™-96 Total RNA Isolation Kit (ThermoFisher AM1830) and quantified (ThermoFisher Q10211); cDNA was generated from 500 ng of RNA (ThermoFisher 11754250). qRT-PCR was performed using the TaqMan Fast Advanced Master Mix (ThermoFisher 4444557) with the FX96 Real-Time PCR Detection System (Bio-Rad) with TaqMan probes (**Table S9**). The results are expressed as fold-increase mRNA expression of the gene of interest normalized to TBP expression by the ΔΔCt method (**Table S10**, some samples omitted from lack of amplification).

Two gRNAs produced an outsized effect on growth inhibition compared to the other gRNAs targeting the same DHS at 1-week post-transduction in the gRNA abundance screen and throughout the growth competition assay (**Fig. 5C**, top left, DHSs chr2.6452 and chr1.10482 in Fig. S24). These gRNAs were detected in a substantially greater number of cells in the scRNA-seq screen but visibly depleted the cell population during culture and impacted cell fitness enough within the first week that we were unable to collect sufficient quantities of quality RNA to perform qPCR amplification; these DHS-gene pairs are not displayed in qPCR analysis. These gRNAs could be individually toxic or have a much larger effect on target gene expression than other gRNAs in the DHS, but their rapid effect on cell fitness makes their gene expression effects difficult to measure at the screen time point.

For growth competition assays, 3×10^4^ cells were transduced with lentivirus encoding a single gRNA and GFP into polyclonal K562 dCas9^KRAB^ cells. Cells were transduced with either 1) an individual targeting gRNA and GFP or 2) non-targeting gRNA and GFP or 3) a pool of all (4-5) gRNAs targeting a particular DHS in the screen. After 2 days, transduced cells were mixed 1:1 with untransduced cells of the same polyclonal line. Competition assays were seeded in 5 replicates per gRNA at ∼10k cells per well in a 96-well plate. Plates were observed in the Incucyte for 15 days with phase and GFP imaging GFP every 6 hours. Growth is expressed as the ratio of green area to total phase area, relative to the beginning of the Incucyte observation (day 0 in **Table S11**).

## List of Supplementary Tables

|  |  |
| --- | --- |
| Table S1 | original wgCERES discovery cell fitness screen gRNA information, raw counts and DESeq2 results. Duplicates removed. |
| Table S2 | Bulk DHS summary results table. Duplicates removed. |
| Table S3 | K562 validation cell fitness screen gRNA information, raw counts and DESeq2 results. Duplicates removed. |
| Table S4 | OCIAML2 cell fitness screen gRNA information, raw counts and DESeq2 results. Duplicates removed. |
| Table S5 | gRNAs used for single-cell experiment, with raw counts and DESeq2 results for 1-week cell fitness screen. |
| Table S6 | 1-week single-cell RNA-seq screen results table (individual gRNA-gene pairs). |
| Table S7 | 1-week single-cell RNA-seq screen results (aggregated to DHS-gene pairs). |
| Table S8 | Individual validation gRNAs used for qRT-PCR and competition assays. |
| Table S9 | Taqman probes for qRT-PCR validation. |
| Table S10 | Individual validation qPCR results. |
| Table S11 | Individual validation competition assay results. |

## Data Availability

All raw and processed sequencing files and Supplementary Tables and Data can be accessed via the Gene Expression Omnibus under accession number GSE167890 with secure token ajefmwoirpyjhsf.

## Acknowledgements

This work was supported by the US National Institutes of Health (NIH) grants to GEC, TER, and CAG (DP2OD008586, R01DA036865, UM1HG009428, R01HG010741, RM1HG011123, U01AI146356, UM1HG012053, R01CA289574, and R01MH125236), an Allen Distinguished Investigator Award from the Paul G. Allen Frontiers Group, the Open Philanthropy Project, and a National Science Foundation (NSF) grant (EFMA-1830957). T.S.K. was supported by a NIH Biotechnology Training Grant (T32GM008555). We thank Alejandro Berrio Escobar and Greg Wray for help with comparative chromatin analysis, and the DCI Flow Core for using the FACSCanto. We thank the Duke University School of Medicine for the use of the Sequencing and Genomic Technologies Shared Resource for sequencing support. This work used a high-performance computing facility partially supported by grants 2016-IDG-1013 (“HARDAC+: Reproducible HPC for Next-generation Genomics“) and 2020-IIG-2109 (“HARDAC-M: Enabling memory-intensive computation for genomics“) from the North Carolina Biotechnology Center. Parts of some figures were created in https://BioRender.com.

