## Supplementary Figures for "Genome-wide annotation of gene regulatory elements linked to cell fitness"

**Figure S1: Overview of gRNA design for whole-genome discovery, validation, and 1-week cell fitness screens.**

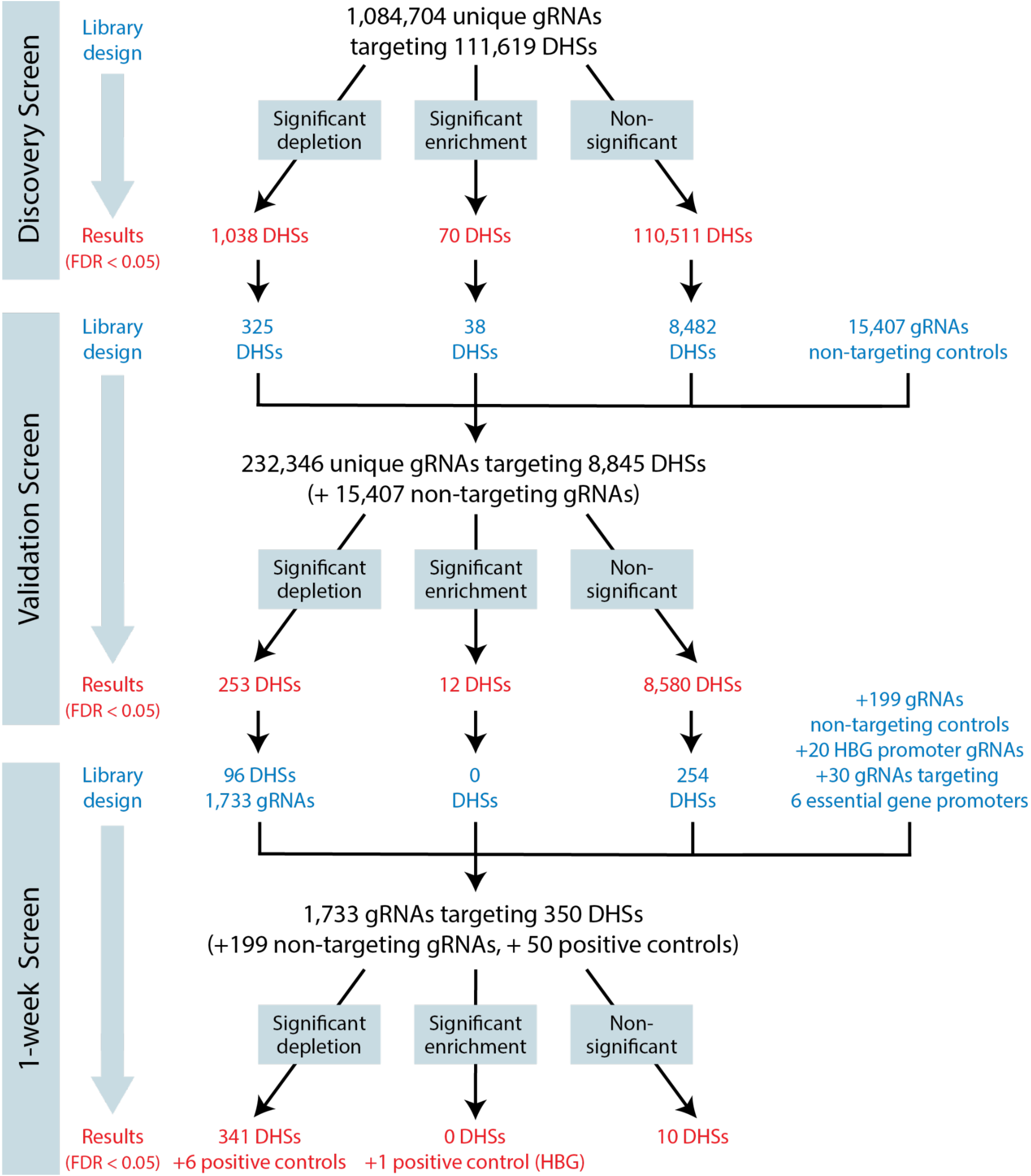


**Figure S1: Overview of gRNA design for whole-genome discovery, validation, and 1-week cell fitness screens.** Shown are the number of tested and control gRNAs that are used for each screen, the number of DHS sites that are detected as significant, and the ones that were included in subsequent screens.

**Figure S2: Characterization of whole-genome discovery screen gRNA library.**

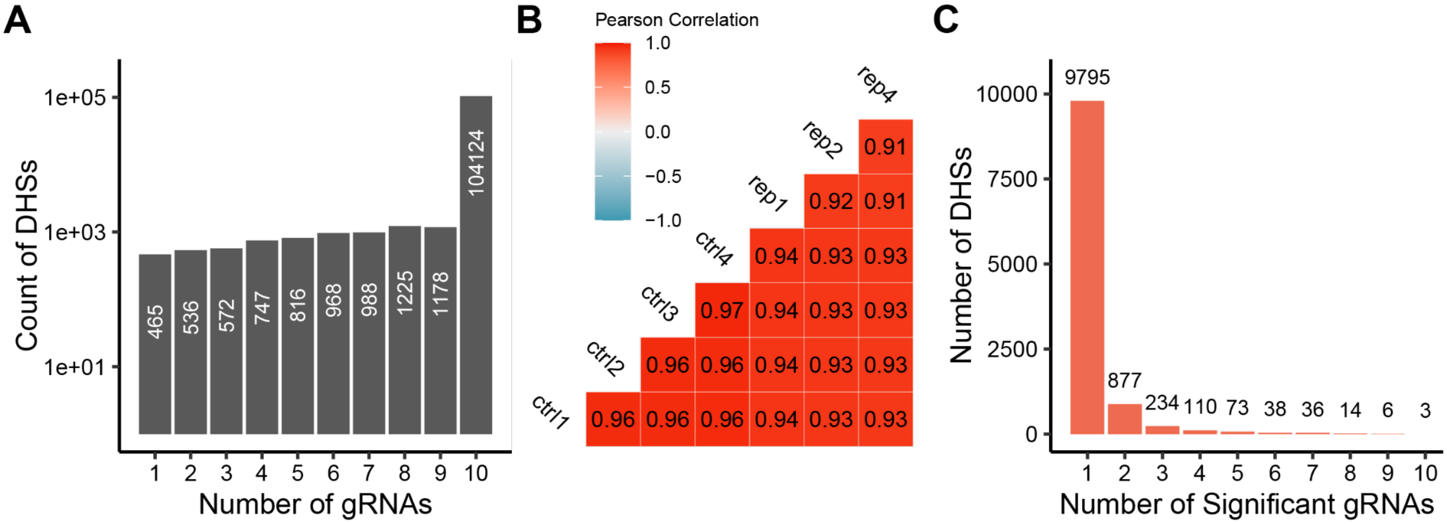


**Figure S2: Characterization of whole-genome discovery screen gRNA library. (A)** Number of gRNAs in each DHS in the discovery screen library. Almost all DHSs had 10 gRNAs. **(B)** Correlation of raw gRNA sequencing read counts across replicates in the genome-wide discovery screen. **(C)** Number of significant gRNAs (FDR < 0.1) per DHS in the genome-wide discovery screen among DHSs with significant gRNAs; most have 1 significant gRNA.

**Figure S3: Non-essential Gene Promoters and Number of Significant gRNAs per DHS**

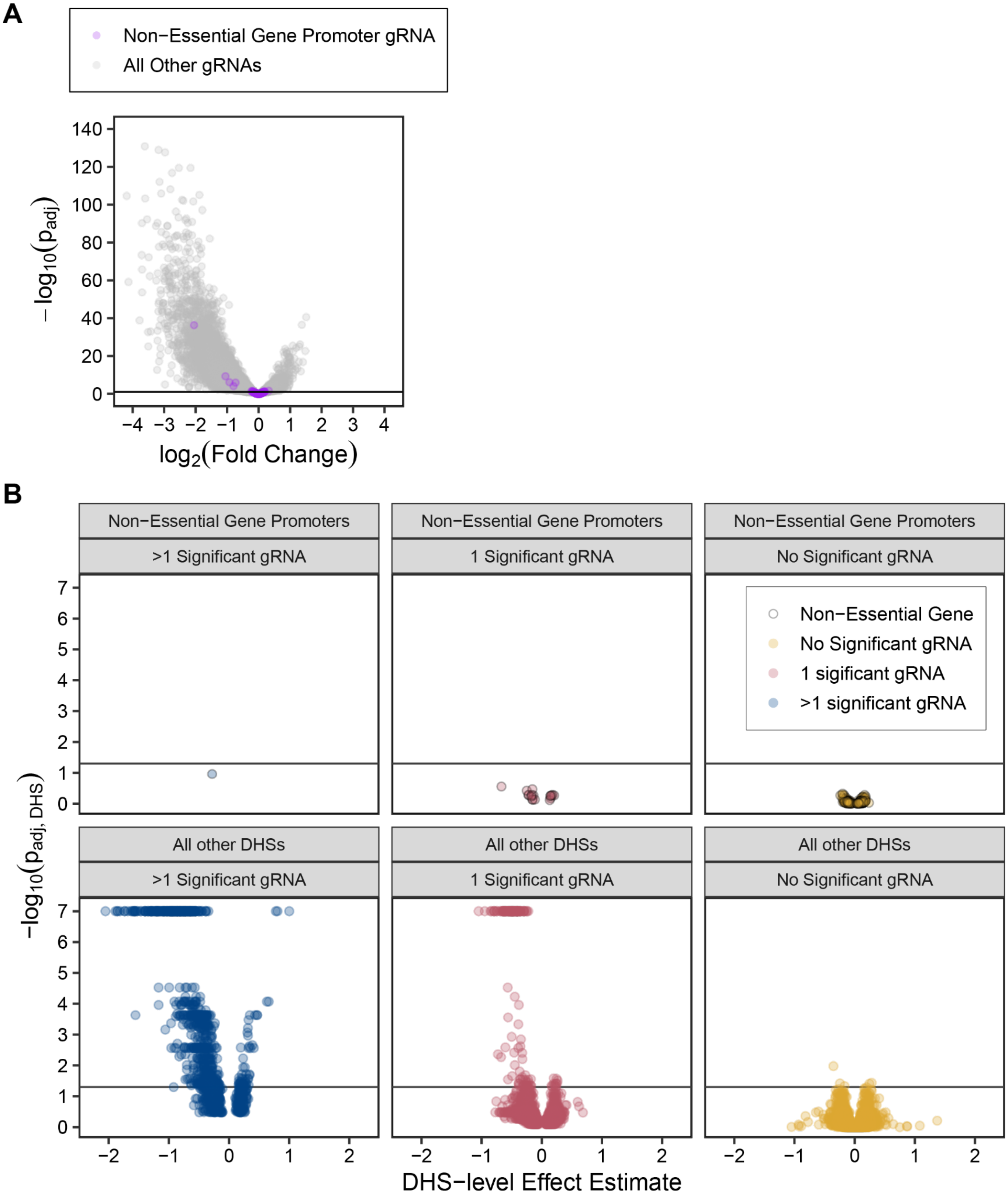


**Figure S3: Non-essential Gene Promoters and Number of Significant gRNAs per DHS. (A)** gRNA-level volcano plot for the discovery screen with gRNAs targeting negative control promoters highlighted in purple. **(B)** DHS-level volcano plot for the discovery screen, divided by whether or not the DHS is a negative control (non-essential gene) promoter and the number of significant gRNAs in the DHS. Of the significant DHSs, 923 had more than 1 significant gRNA, 180 DHSs had 1 significant gRNA, and 5 had no significant gRNAs.

**Figure S4: gRNA attributes.**

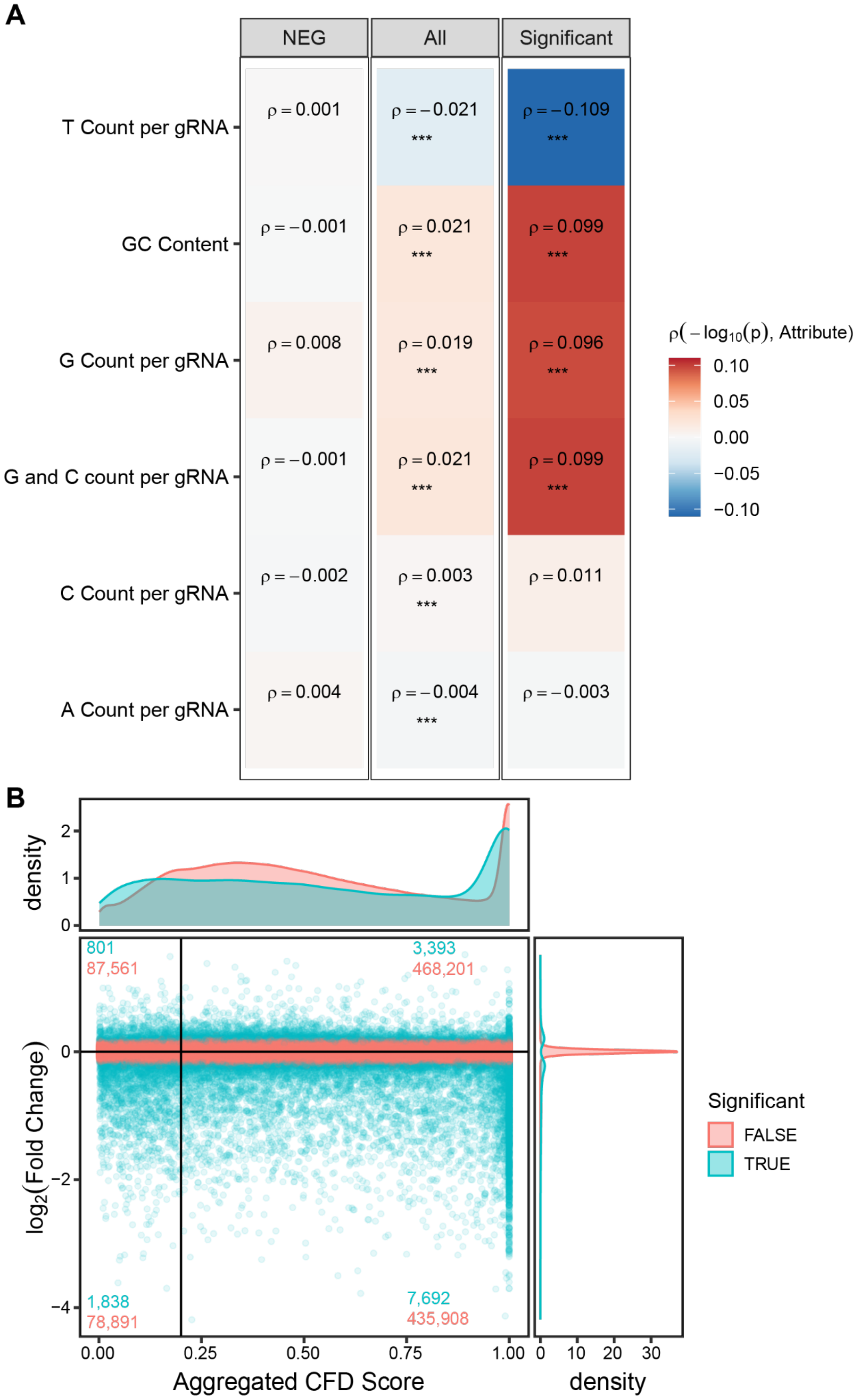


**Figure S4: gRNA attributes. (A)** Correlation of gRNA attributes. Various numerical features were collected gRNAs in nonessential negative control (NEG) DHSs, all DHSs, and significant (FDR < 0.05) DHSs. Displayed are Spearman correlations between individual gRNA significances (expressed as $-log_{10}\, p$ ) and various gRNA attributes in (left) non-essential control DHSs, (middle) all DHSs and (right) only essential DHSs (FDR < 0.05). The Spearman correlation is displayed for each comparison, and significant (p<0.001) Spearman correlation tests are marked with ***. **(B)** Aggregated cutting frequency determination scores (CFD or

Guidescan specificity scores) for all gRNAs in the genome-wide discovery screen, plotted by their enrichment effect size ($log_2\ (FC)$). Counts of both significant and insignificant gRNAs in each quadrant are given on the plot (FDR < 0.1). Significant gRNAs are not enriched in the lower left quadrant (low-specificity gRNAs decreasing cell fitness, p = 0.8141 from a Fisher's exact test).

**Figure S5: Genome-wide regulation of cell fitness in gene bodies, promoters, UTRs, and intergenic regions.**

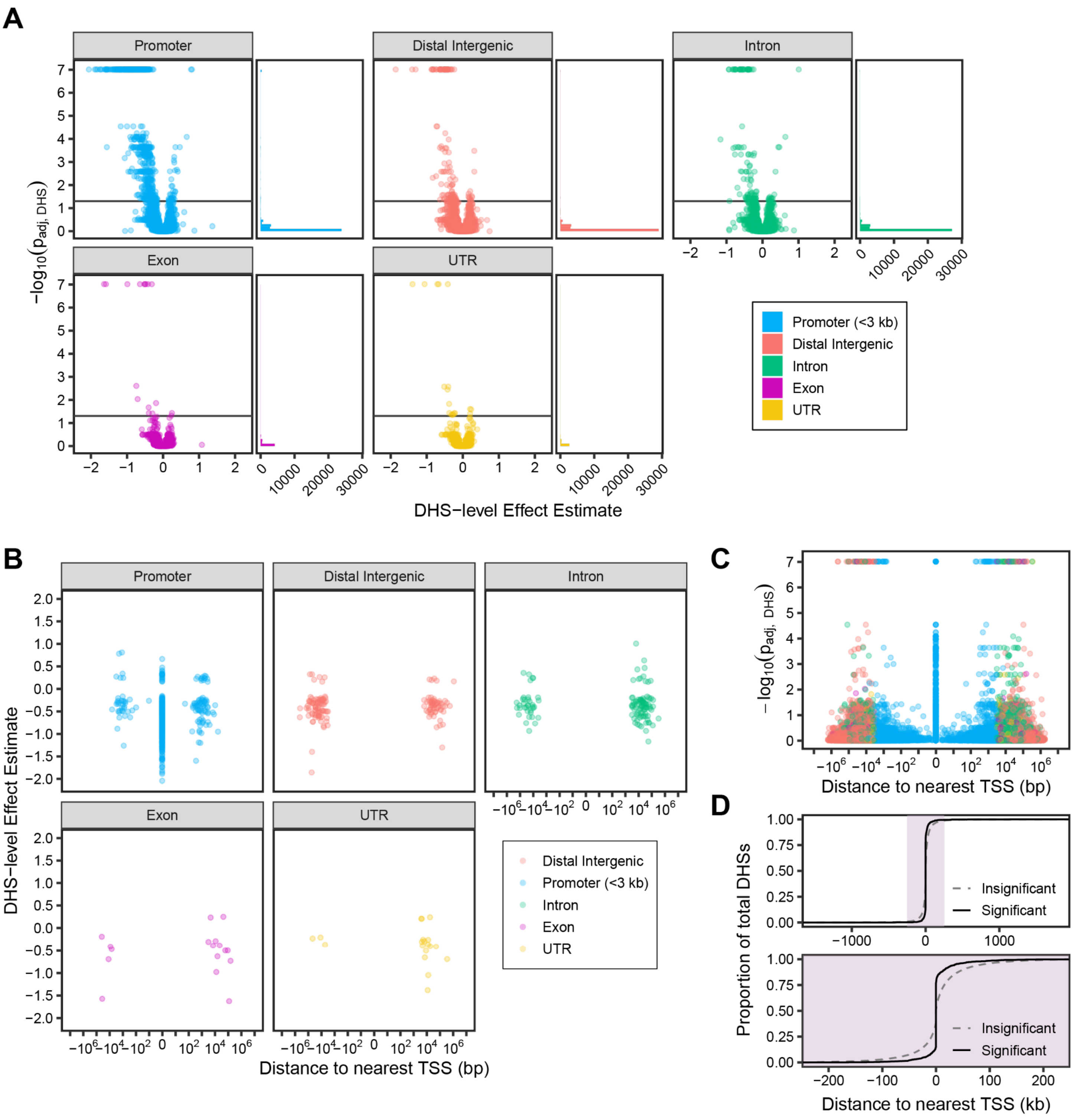


**Figure S5: Genome-wide regulation of cell fitness in gene bodies, promoters, UTRs, and intergenic regions. (A)** DHS-level volcano plot for the discovery screen, divided by region type. **(B)** Distribution of hit DHS effect size estimates by their distance from the nearest transcription start site (TSS), divided by region type. **(C)** Distribution of DHS-level p-values by the distance of the DHS from the nearest transcription start (TSS). **(D)** ECDFs showing the position of significant and insignificant DHSs relative to the nearest TSS genome-wide. Top: full library, Bottom: Same as (Top) but with the scale restricted to +/- 250 kb.

**Figure S6: Distal Lmo2 enhancers linked to cell fitness.**

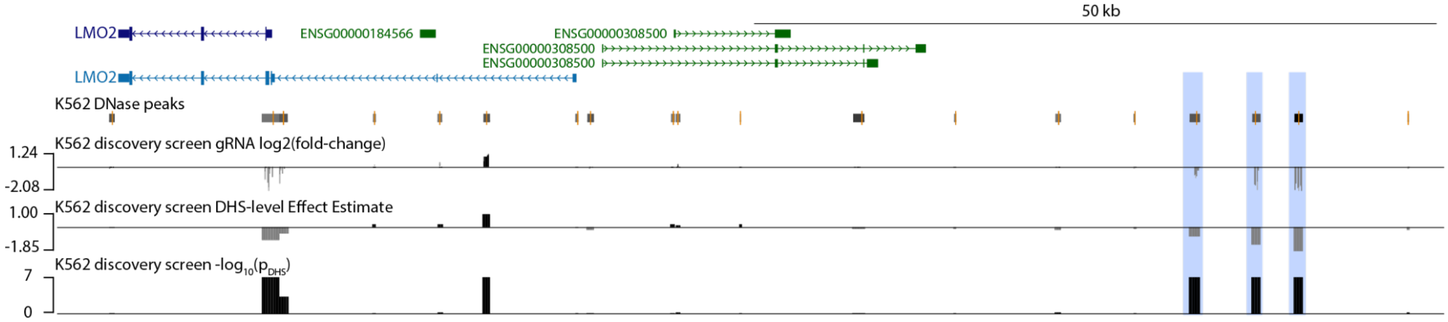


**Figure S6: Distal Lmo2 enhancers linked to cell fitness.** Genome browser tracks of three depleted DHSs upstream of *Lmo2* (highlighted regions) and 3 significant DHSs in the gene (hg19 chr11:33,875,577-33,977,115).

**Figure S7: Correlation analysis of DHS attributes**

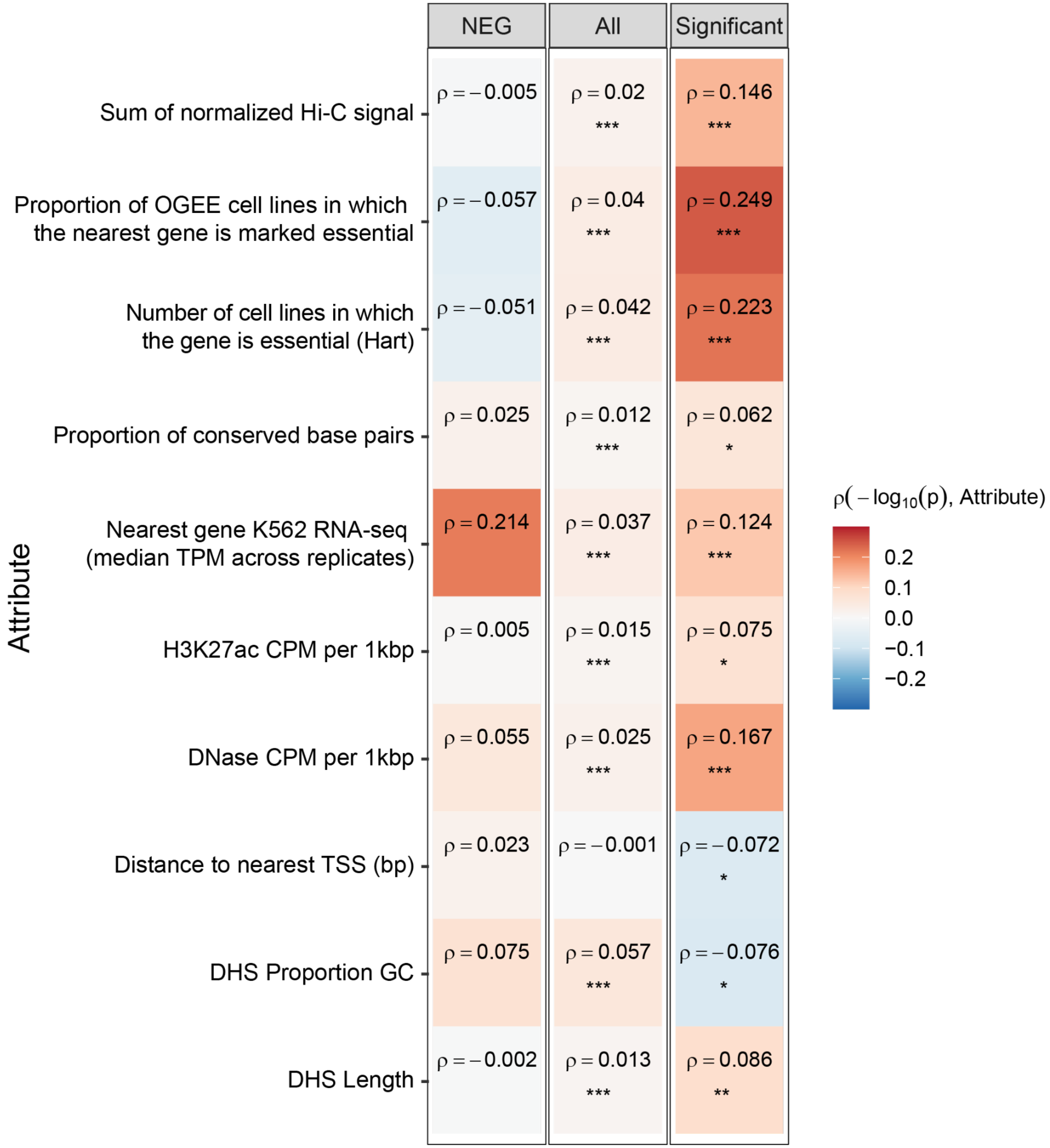


**Figure S7: Correlation analysis of DHS attributes**. Various numerical features were collected for negative control DHSs (left column), all DHSs (middle), and significant DHSs (right, FDR < 0.05). Displayed is the Spearman correlation between the DHS significance score from the RRA test, expressed as $-\log_{10} p$ , and the numerical value of each DHS attribute. The Spearman correlation is displayed for each comparison, and significant Spearman correlation tests are marked: *** p< 0.001, ** p < 0.01, * p< 0.05.

**Figure S8: Comparison of promoter wgCERES DHS hits to other published essentiality studies**

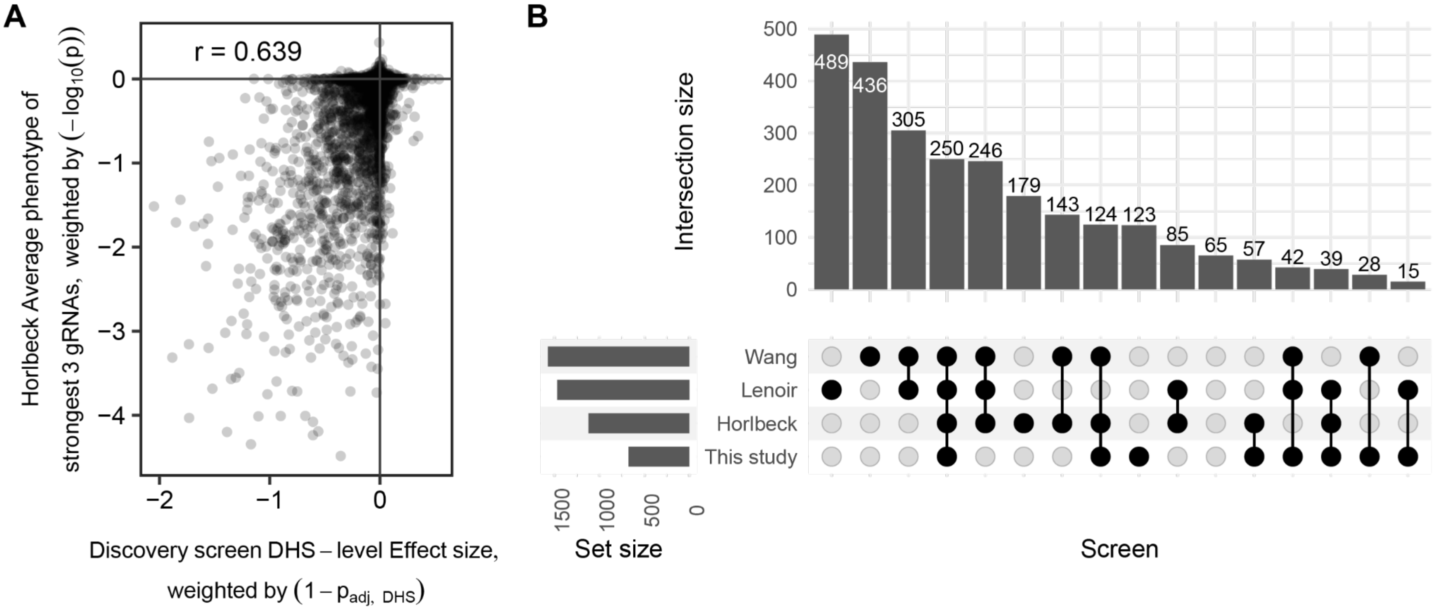


**Figure S8: Comparison of promoter wgCERES DHS hits to other published essentiality studies.** Three previous studies have performed essentiality studies in K562 cells, including one study that performed CRISPRi targeting promoters (Horlbeck et al. 2016) and 2 studies using CRISPR/Cas9 targeting exons (Lenoir et al. 2018; Wang et al. 2017). **(A)** Gene-level comparison of weighted effect sizes of 18,931 gene promoters in this study with growth phenotype from promoter-targeting CRISPRi (y-axis). These results compare the average across promoter annotations from the Horlbeck dataset to the single DHS from this dataset closest to the gene promoter. Note that there is a general correspondence between assays. **(B)** “UpSet” plot showing overlap of significant promoter/gene hits identified by all 4 studies. Note that in addition to these comparisons, this study identified and characterized distal non-promoter regulatory elements not identified by previous studies.

**Figure S9: Validation Screen gRNA-library characterization and hits per DHS**

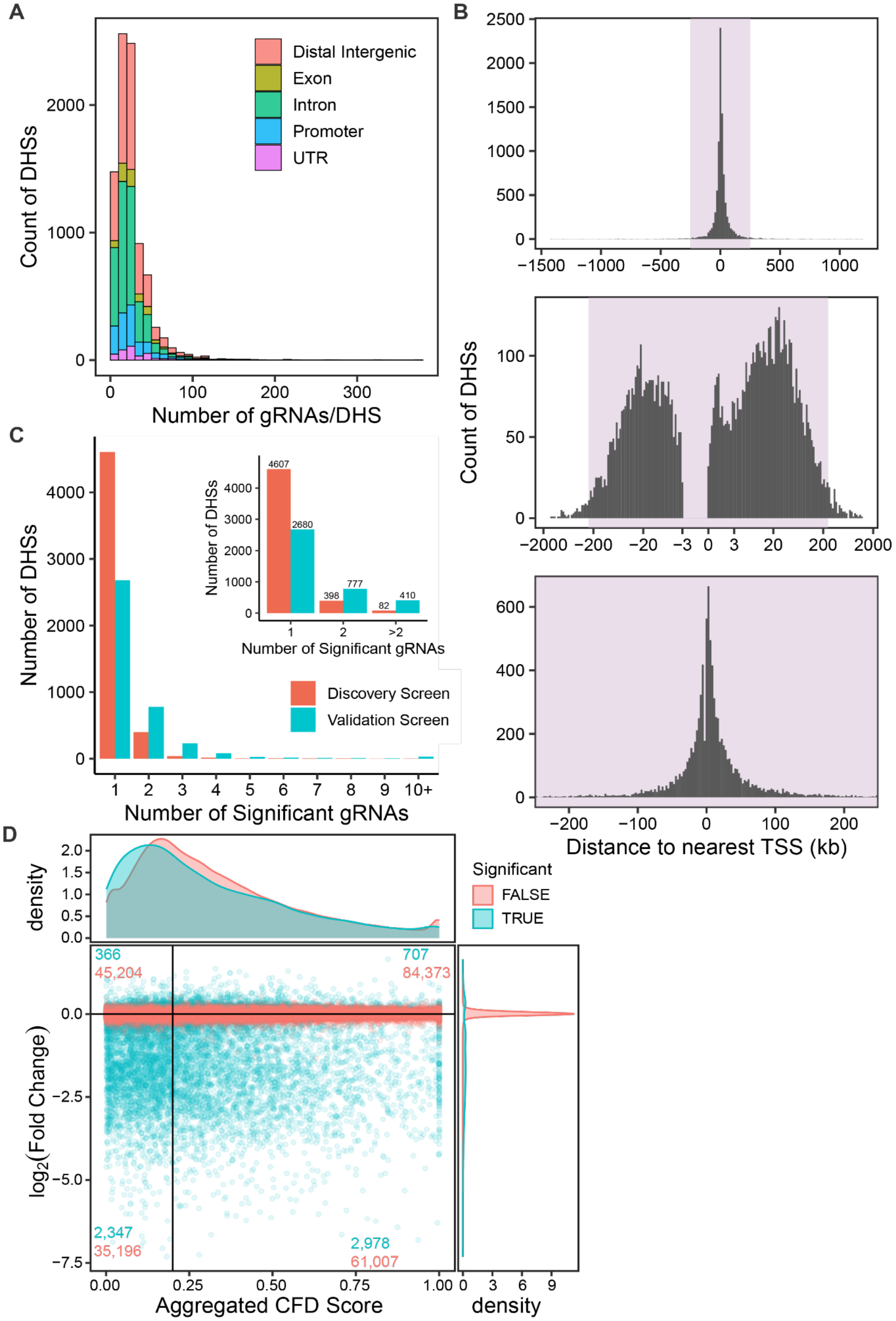


**Figure S9: Validation Screen gRNA-library characterization and hits per DHS. (A)** Distribution of the number of gRNAs per DHS in the validation screen library, colored by the type of region. **(B)** Distribution of distance to nearest transcription start site (TSS) in the validation screen library. Top: full library, Middle: log-scaled full library showing the exclusion of most promoters just upstream of the TSS, Bottom: Same as (Top) but with the scale restricted to +/- 250 kb. **(C)** Counts of the number of significant gRNAs per DHS in the K562 validation and discovery screens between the shared 8,833 DHSs with data in both screens. **(D)** Aggregated cutting frequency determination scores (CFD or Guidescan specificity scores) for gRNAs in the distal sublibrary screen in K562s, plotted by their enrichment effect size ($log_2\ (FC)$). Counts of both significant and insignificant gRNAs (FDR = 0.05) in each quadrant are given on the plot. Significant gRNAs are enriched in the lower left quadrant (low-specificity gRNAs decreasing cell fitness, p < 0.00001 from a Fisher's exact test).

**Figure S10: Validation Screen DHS Volcano Plots.**

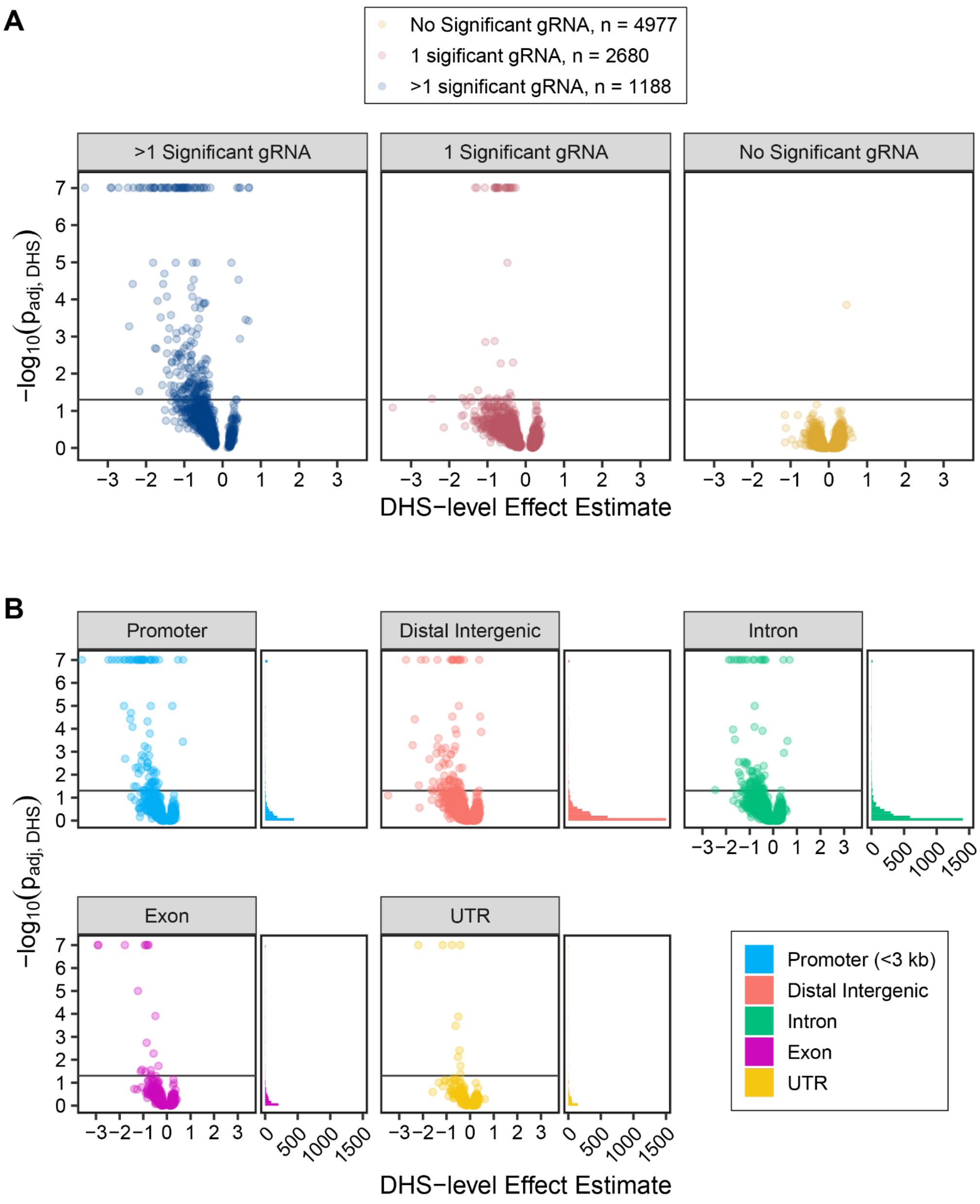


**Figure S10: Validation Screen DHS Volcano Plots. (A)** Volcano plot of significance of DHS' effect on cell growth in K562s relative to aggregate effect size estimate, colored and divided by the number of individually significant gRNAs in the DHS (FDR < 0.05). A DHS-level false discovery rate of 0.05 is marked with a horizontal line. Of the significant DHSs, 227 had more than 1 significant gRNA, 37 had 1 significant gRNA, and 1 had no significant gRNAs. **(B)** Volcano plot of significance of DHS' effect on cell growth in K562s relative to aggregate effect size estimate, divided and colored by the type of region. A DHS-level false discovery rate of 0.05 is marked with a horizontal line across all plots.

**Figure S11: Effects of both individual gRNAs and aggregated DHSs correlate between Discovery and Validation Screens**

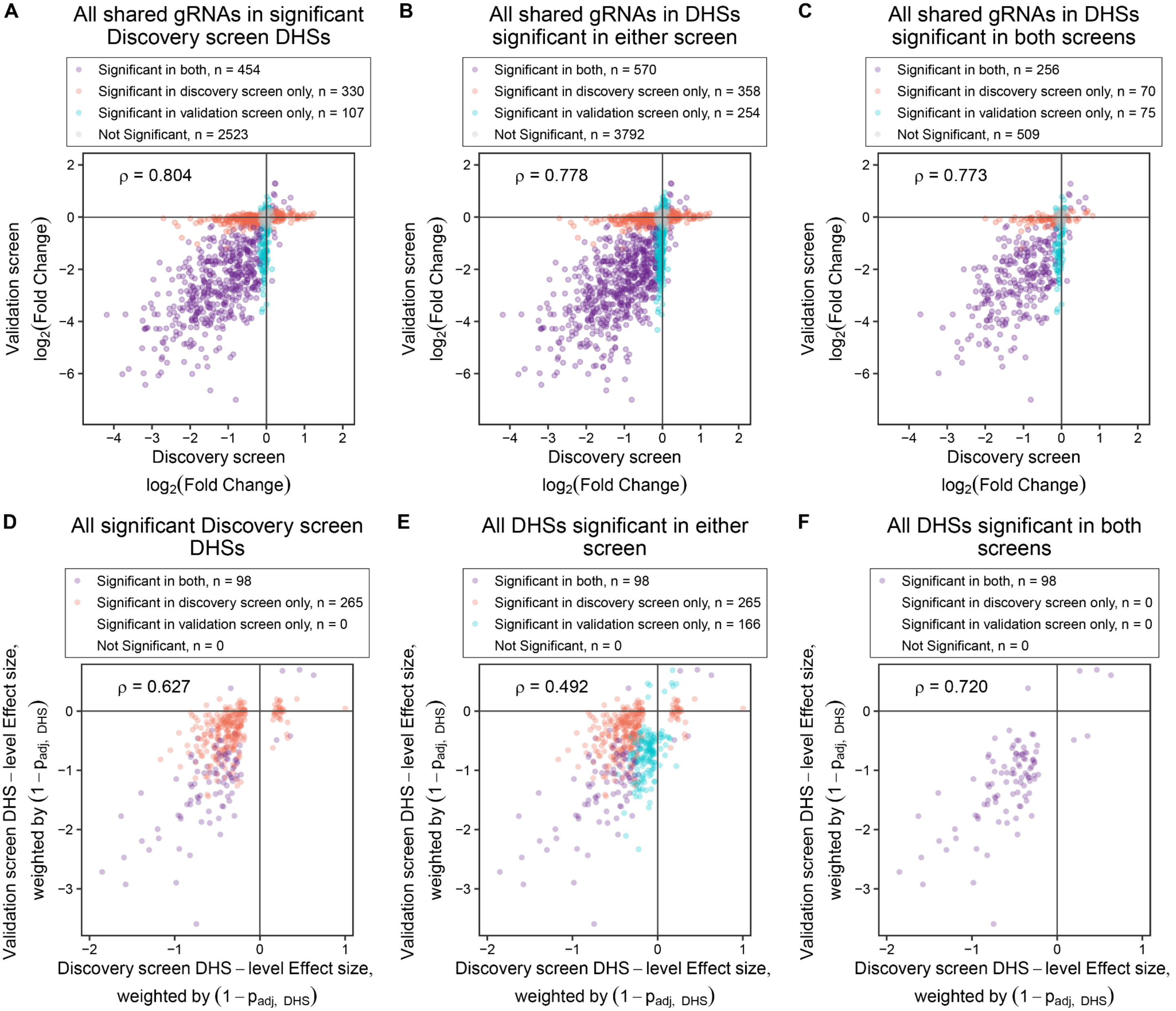


**Figure S11: Effects of both individual gRNAs and aggregated DHSs correlate between Discovery and Validation Screens. (A-C)** Correlations between fold changes in gRNA abundance in the initial discovery screen and distal sublibrary screen across all protospacers shared between both screens, in **(A)** all DHSs called significant (FDR < 0.05) in the genome-wide discovery screen, **(B)** the union set of DHSs significant (FDR < 0.05) in either screen and **(C)** the intersection set of DHSs significant (FDR < 0.05) in both screens. **(D-F)** Correlations between weighted DHS-level effect size estimates in the initial discovery screen and distal sublibrary screen in **(D)** all DHSs called significant (FDR < 0.05) in the genome-wide discovery screen, **(E)** the union set of DHSs significant (FDR < 0.05) in either screen and **(F)** the intersection set of DHSs significant (FDR < 0.05) in both screens.

**Figure S12: Sensitivity, Specificity, and Precision as functions of gRNA and DHS-level FDRs.**

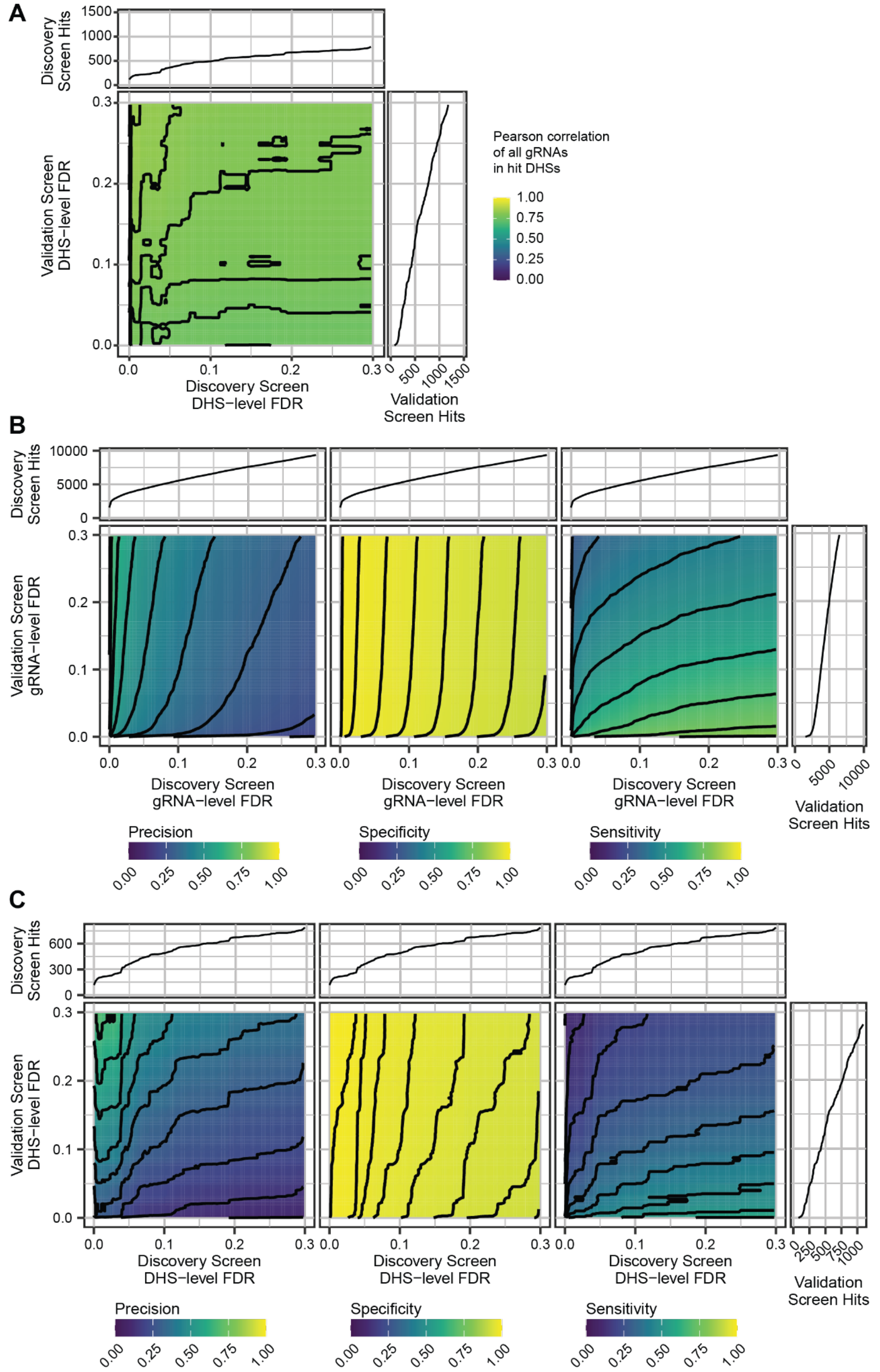


**Figure S12: Sensitivity, Specificity, and Precision as functions of gRNA and DHS-level FDRs. (A)** Heatmap of Pearson correlation between gRNA enrichment effect sizes ($log_2(FC)$) in discovery and validation screens, restricted to gRNAs in DHSs significant at the given FDR on the x and y axes. Notably, the correlation is highly robust to adjustments in DHS-level FDR. **(B-C)** Analysis of (left) precision, (middle) specificity, and (right) sensitivity of the initial discovery screen as a function of **(B)** gRNA-level or **(C)** DHS-level false discovery rates, using the validation screen results as ground truth. The marginal plots indicate accumulation of individually significant gRNAs in either screen as a function of the **(B)** gRNA-level or **(C)** DHS-level false discovery rate (FDR).

**Figure S13: Comparison of replicate correlations in K562 and OCI-AML2 validation screens**

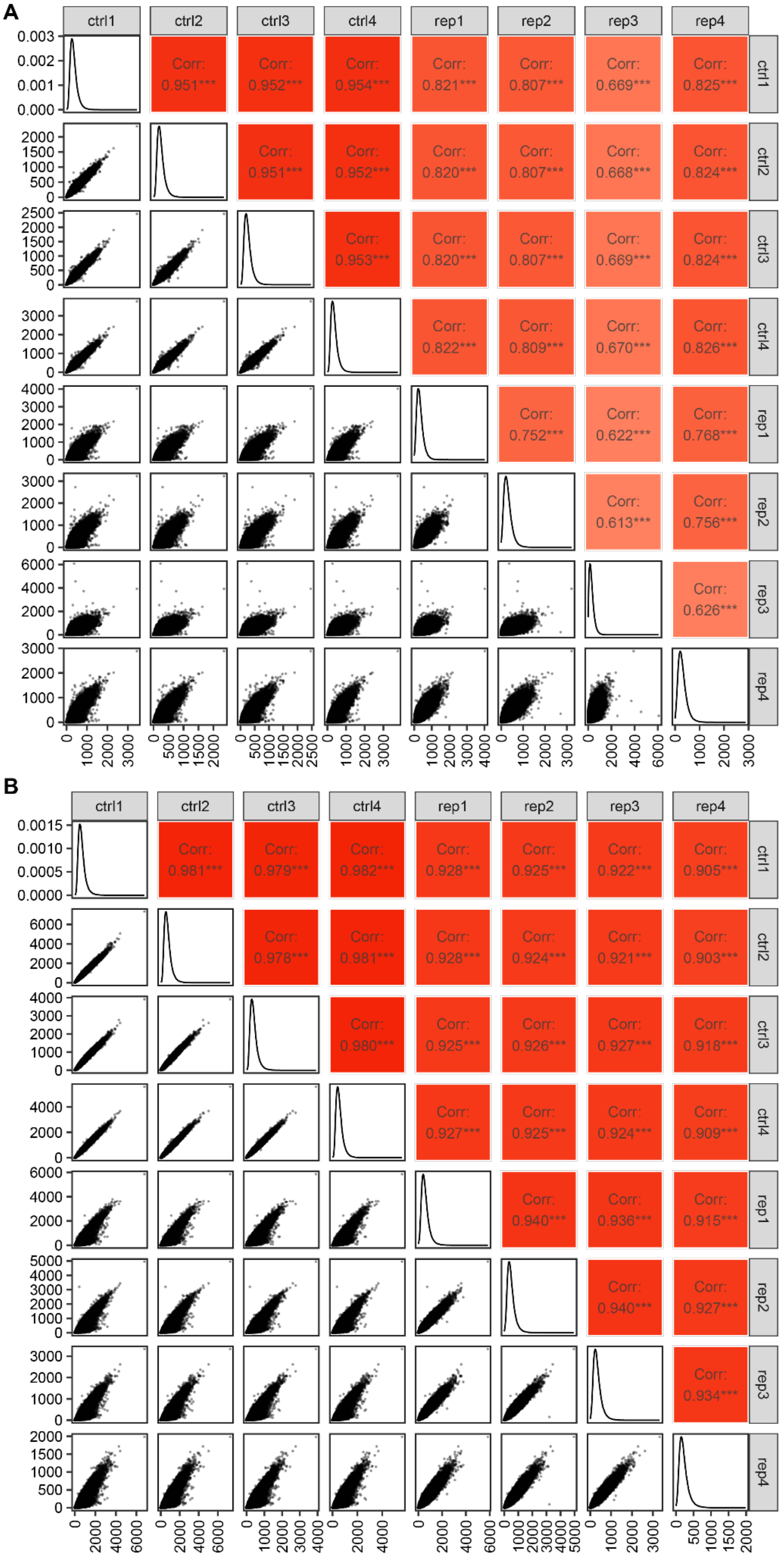


**Figure S13: Comparison of replicate correlations in K562 and OCI-AML2 validation screens.** Replicate correlations (Pearson's r) of raw gRNA read counts in the distal sublibrary validation screen in **(A)** K562 cells and **(B)** OCI-AML2 cells. Each point represents a different protospacer sequence.

**Figure S14: OCI-AML2 fitness screen volcano plots.**

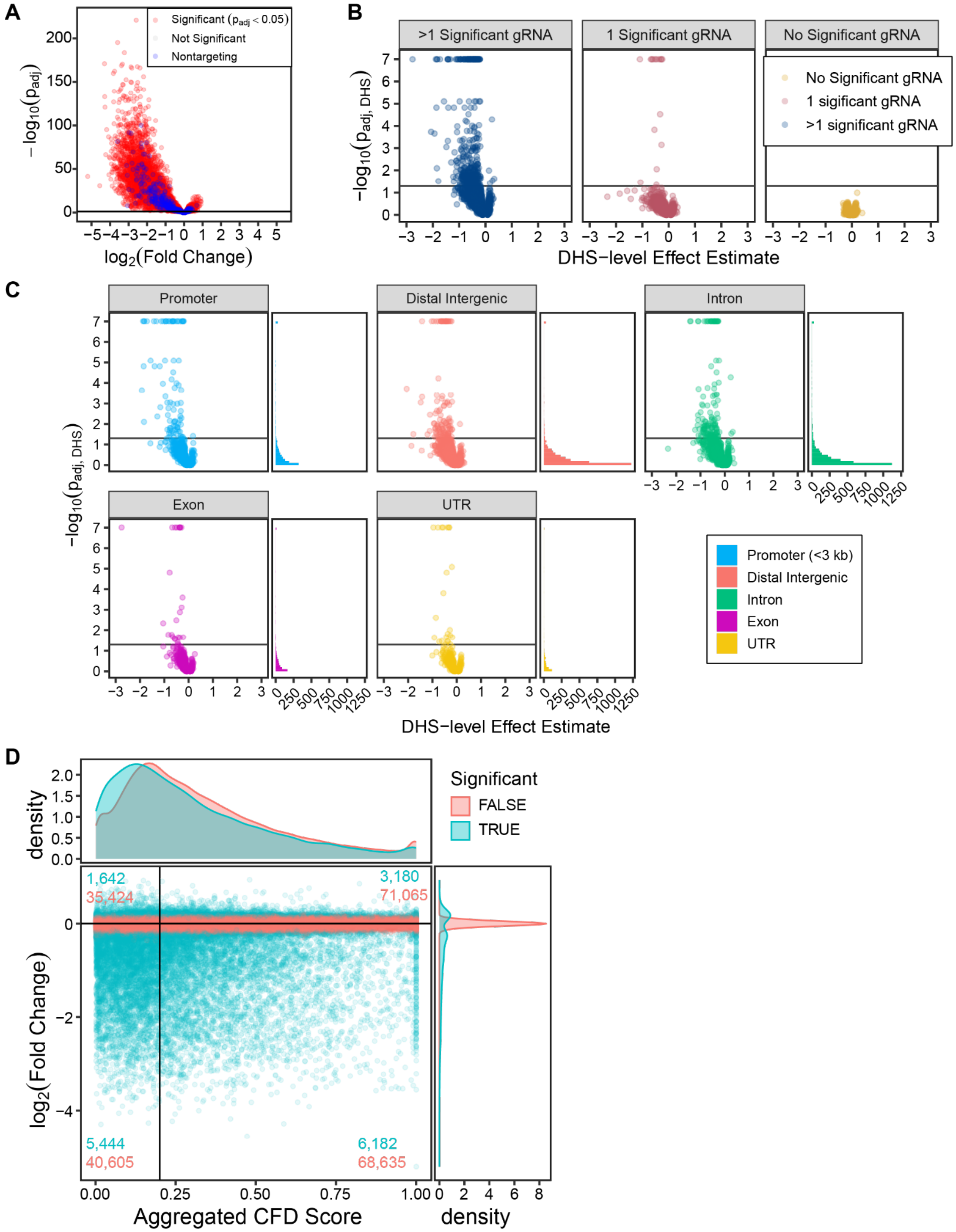


**Figure S14: OCI-AML2 fitness screen volcano plots. (A)** gRNA-level volcano plot from the validation screen in OCI-AML2 cells. **(B)** DHS-level volcano plot from the validation screen in OCI-AMl2 cells, divided by the number of significant gRNAs per DHS. **(C)** DHS-level volcano plot from the validation screen in OCI-AMl2 cells, divided by the region annotation. **(D)** Aggregated cutting frequency determination scores (CFD or Guidescan specificity scores) for gRNAs in the distal sublibrary screen in OCI-AML2s, plotted by their enrichment effect size ($log_2\ (FC)$). Counts of both significant and insignificant gRNAs (FDR = 0.05) in each quadrant are given on the plot. Significant gRNAs are enriched in the lower left quadrant (low-specificity gRNAs decreasing cell fitness, p < 0.00001 from a Fisher's exact test).

**Figure S15: K562 and OCI-AML2 cell fitness effects correlate more strongly in shared essential elements.**

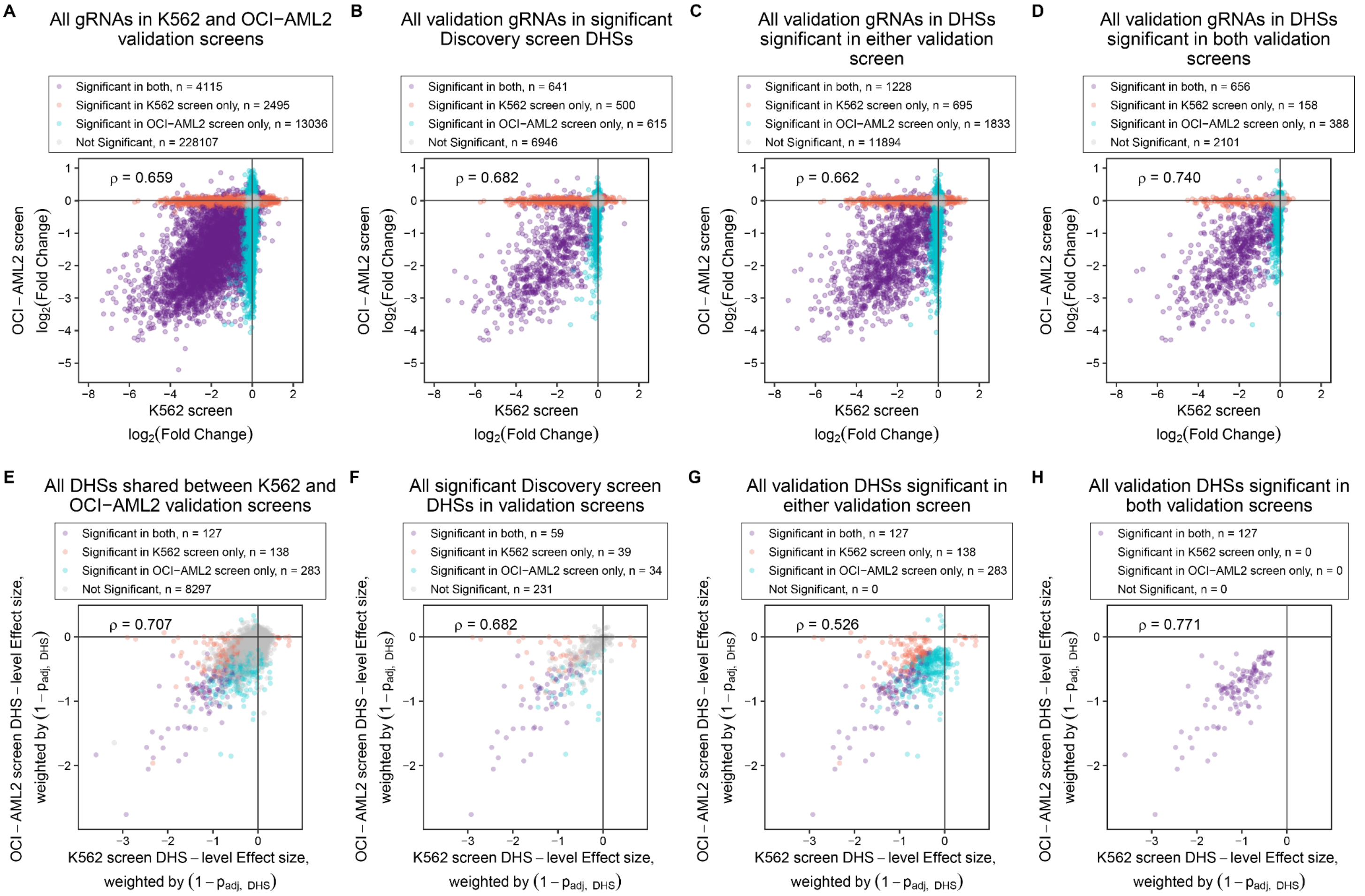


**Figure S15: K562 and OCI-AML2 cell fitness effects correlate more strongly in shared essential elements.** (**A-D**) Correlations between effect sizes (log2 fold change in gRNA abundance) in the distal sublibrary screen in K562 and OCI-AML2 cells. (**E-H**) Correlations between weighted DHS-level effect size estimates in the distal sublibrary screen in K562 and OCI-AML2 cells. From left to right, the plots restrict which DHSs' data are plotted, restricting from all DHSs to DHSs called significant in both screens (FDR < 0.05).

**Figure S16: One-week screen library.**

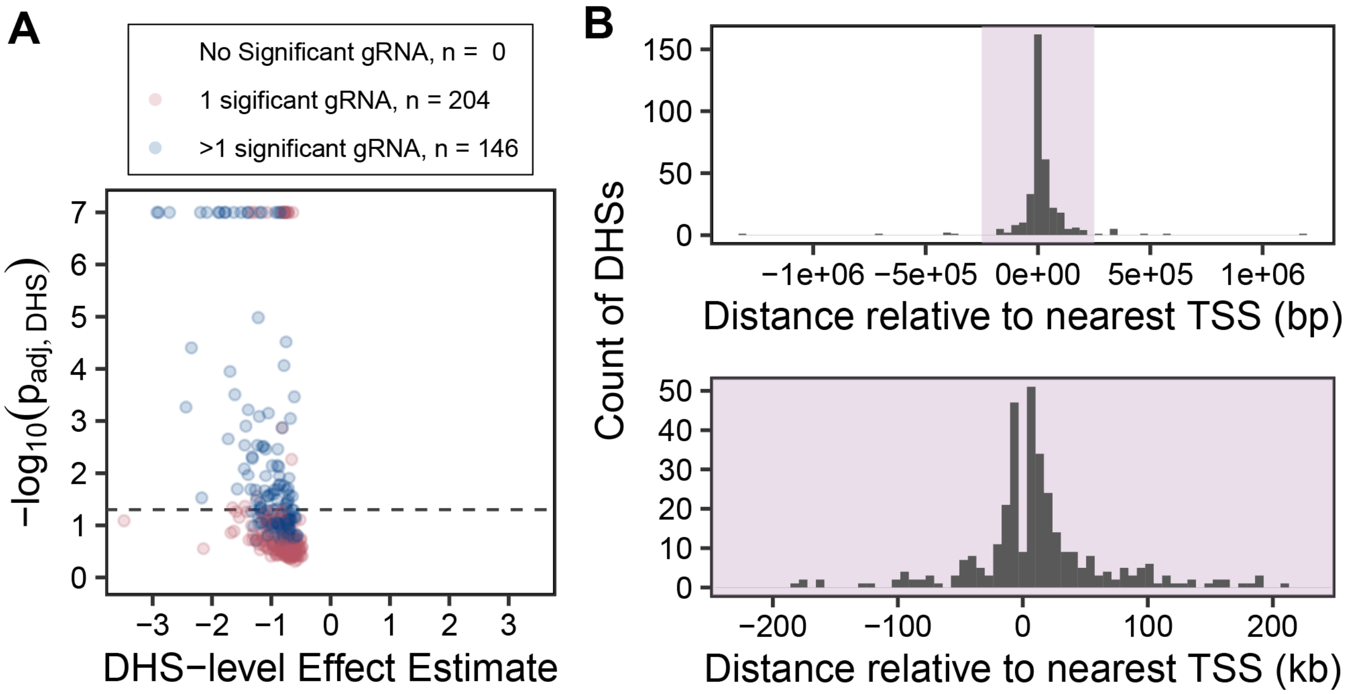


**Figure S16: One-week screen library. (A)** DHS-level volcano plot from the validation screen, subset to the DHSs put into the 1-week scRNA-seq screen. **(B)** Distribution of DHS locations relative to the nearest TSS, for the DHSs put in the 1-week scRNA-seq screen.

**Figure S17: scRNA-seq screen MOI, coverage, and gRNA-gene pair volcano plots.**

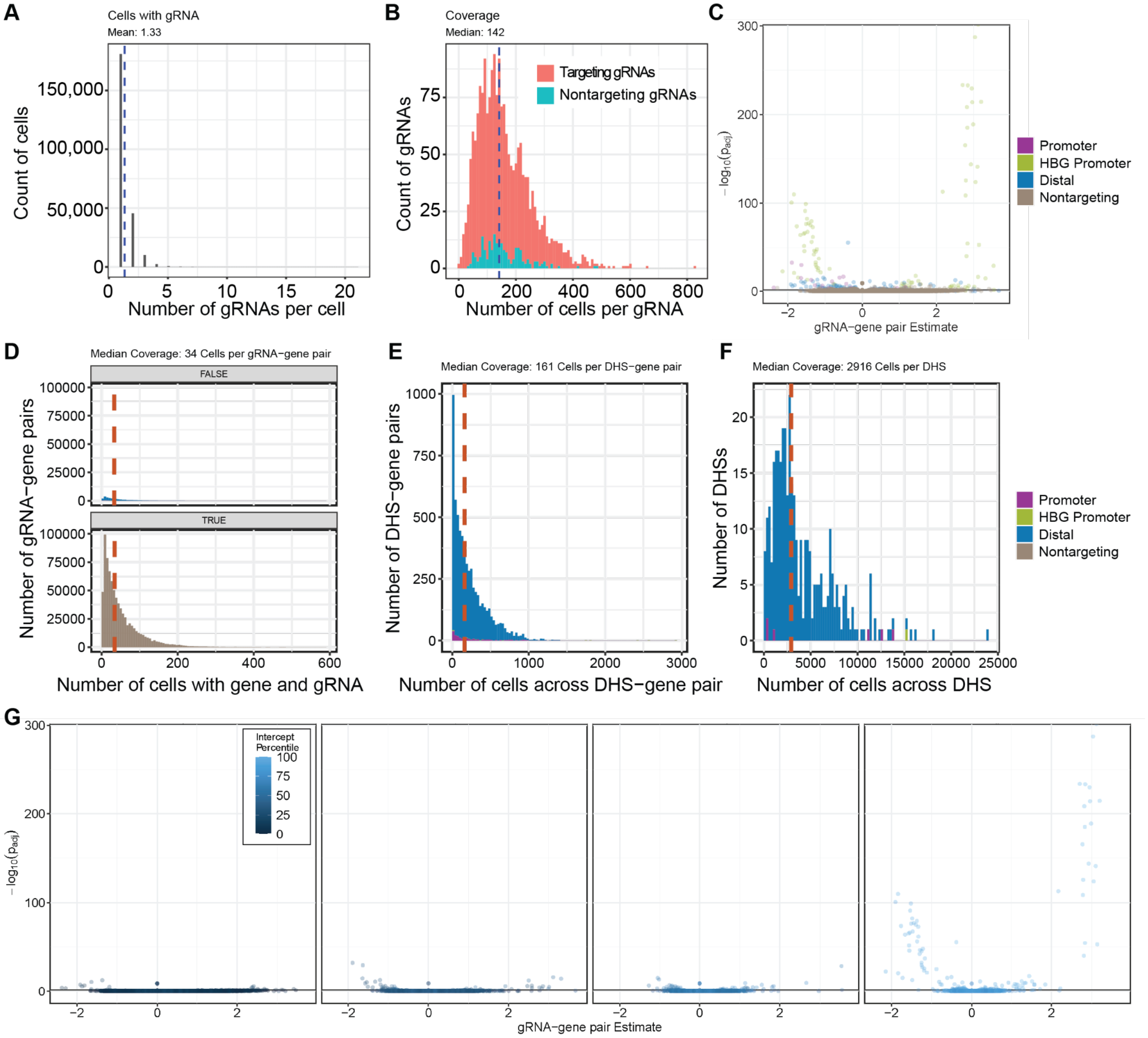


**Figure S17: scRNA-seq screen MOI and coverage. (A)** Histogram of the distribution of the number of gRNAs per cell in the single-cell RNA-seq screen. **(B)** Histogram of the distribution of the number of cells per gRNA (coverage) in the single-cell RNA-seq screen. The blue dashed line marks the median of 142 cells per gRNA. **(C)** Volcano plot of all the gRNA-gene pairs in the single-cell RNA-seq screen, divided by gRNA type. Positive essentiality controls targeting common essential gene promoters are labeled “promoter,” gene expression control gRNAs targeting the HBG promoter are marked “HBG promoter,” experimental gRNAs targeting distal regulatory elements are marked “Distal” and nontargeting gRNAs are marked “Nontargeting.” (D) Coverage at the gRNA-gene pair level for the low MOI perturb-seq screen, dividing the non-targeting gRNA-gene pairs from the targeting gRNA-gene pairs. This is the effective sample size for the negative binomial model. Red dashed line marks the median of 34 cells per gRNA across the whole distribution. **(D)** Coverage at the DHS-gene pair level for the low MOI perturb-seq screen. The red dashed line marks the median of 161 cells per DHS-gene pair. **(E)** Coverage at the DHS-level in the low MOI perturb-seq screen. The red dashed line marks the median of 2,916 cells per DHS. **(F)** gRNA-gene pair results, faceted and colored by the negative binomial model intercept, roughly analogous to gene expression. The most significant results come from the most highly expressed genes.

**Figure S18: Comparison of single-cell RNA-seq screen with published essentiality datasets.**

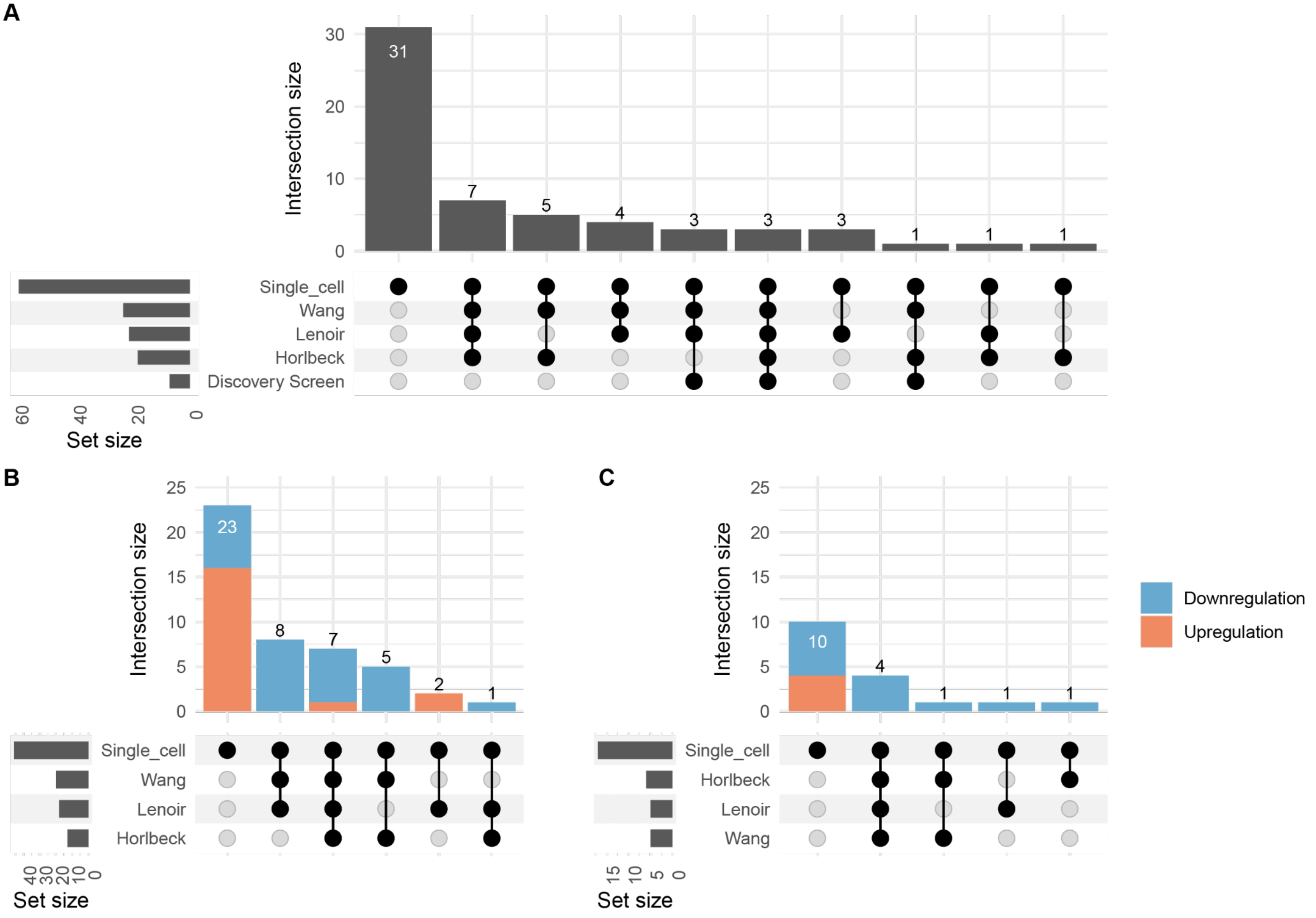


**Figure S18: Comparison of single-cell RNA-seq screen with published essentiality datasets. (A)** Upset plot showing the overlap of 59 unique genes significantly linked to DHSs in this screen with other essential genesets, including one study targeting promoters with CRISPRi (Horlbeck et al. 2016), two studies targeting exons (Wang et al. 2017; Lenoir et al. 2018), and the shared genes from the initial genome-wide discovery screen in this study. Only genes within significant links in this single-cell screen are included on the plot. **(B)** Upset plot showing the overlap of the most significant DHS-gene links per DHS identified in the single-cell screen (46 links total) with previous essentiality screens. **(C)** Upset plot showing the overlap of other significant DHS-gene links per DHS (not the most significant link) identified in the single-cell screen (17 links total).

**Figure S19: Gene regulation qPCR validation results, one week post-transduction.**

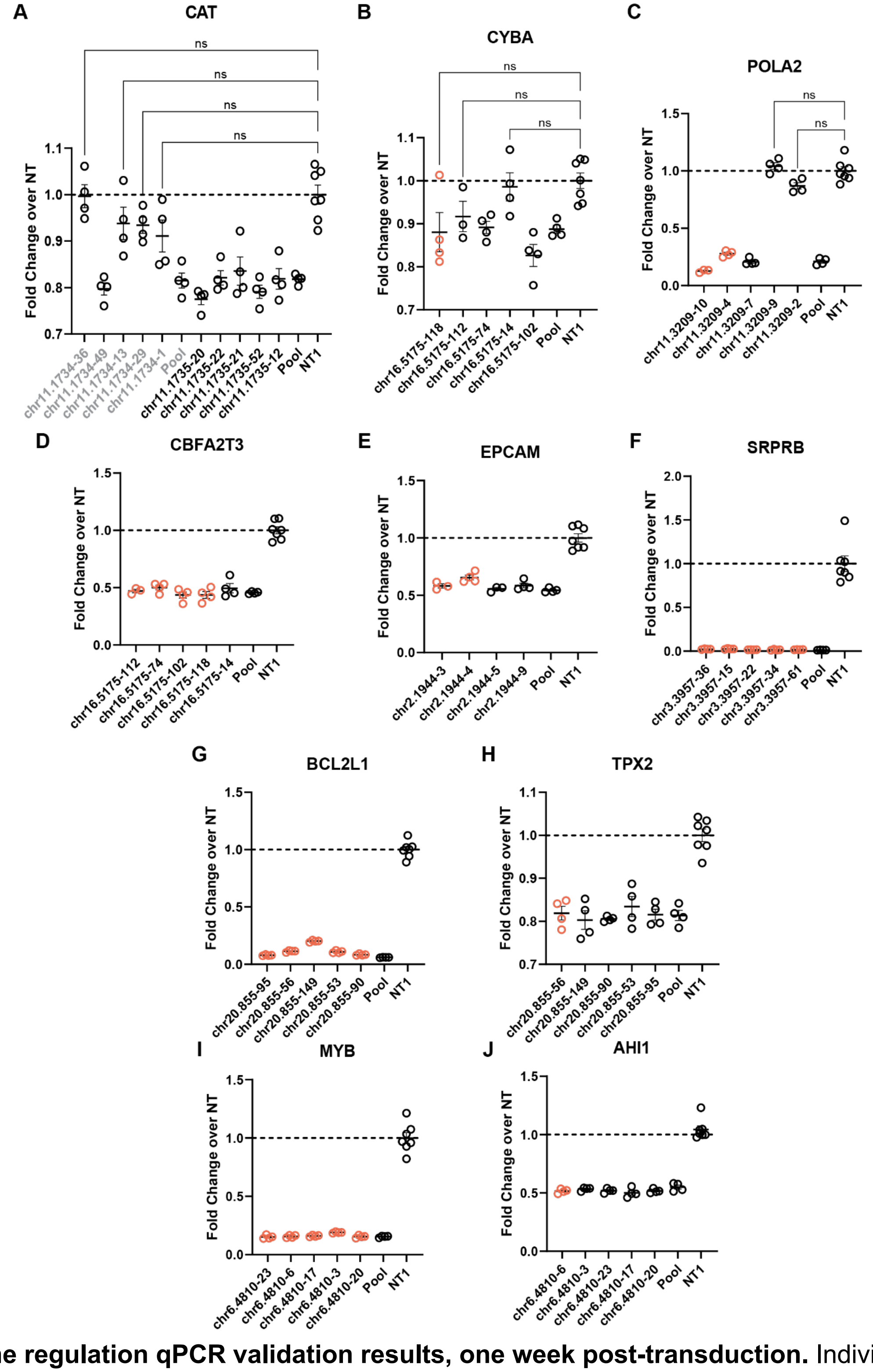


**Figure S19: Gene regulation qPCR validation results, one week post-transduction.** Individual gRNA perturbations' qPCR results are separated by gene and displayed as individual points superimposed on the mean of 4 biological replicates +/- SEM. All DHS-gene pairs validated were significant in the scRNA-seq screen (FDR < 0.05), with the exception of the chr11.1734-CAT gene pair (grey text). Individually significant gRNA-gene links (FDR < 0.05) from the 1-week scRNA-seq screen are displayed with red points, insignificant screen links are colored black. Insignificant differences are marked with ns, otherwise all displayed qPCR data is significant repression in comparison to the nontargeting control (NT1), FDR < 0.05 from Dunnett's T3 multiple comparisons test and Welch ANOVA. Three DHSs validated here were linked to 2 genes, displayed in panels **(B,D)**, **(G-H)** and **(I-J)**.

**Figure S20: Variation within and between gRNAs' effects on target gene expression**

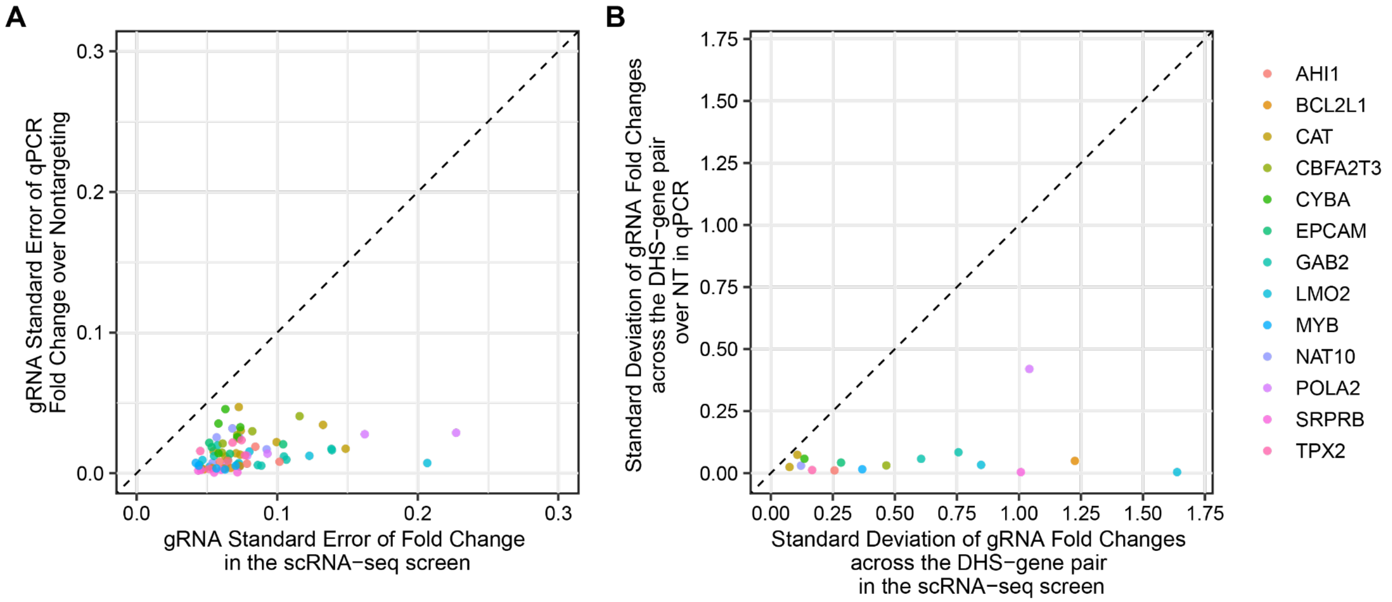


**Figure S20: Variation Within and Between gRNAs' effects on target gene expression. (A)** Comparison of gRNA standard errors in the single-cell RNA-seq screen (median coverage of 142 cells per gRNA) and qPCR (4 biological replicates per gRNA) across the 47 gRNAs and 13 genes with qPCR data in **Fig. 4A**. The dashed line is y = x, representing equal standard errors between the screen and qPCR. **(B)** Comparison of the variability between gRNA-gene pairs within a DHS-gene pair, using standard deviation of the mean fold changes in the qPCR and scRNA-seq screen. The dashed line is y = x, representing equal standard deviations between the screen and qPCR. The one DHS-gene pair with higher variability between gRNAs (POLA2-chr11.3209) was the only DHS whose strongest gene link from the screen had individual gRNA perturbations that were insignificant in the qPCR validation (qPCR data in **Fig. S19C**).

**Figure S21: Transcriptome-Wide Differential Expression Analysis of Validation DHSs.**

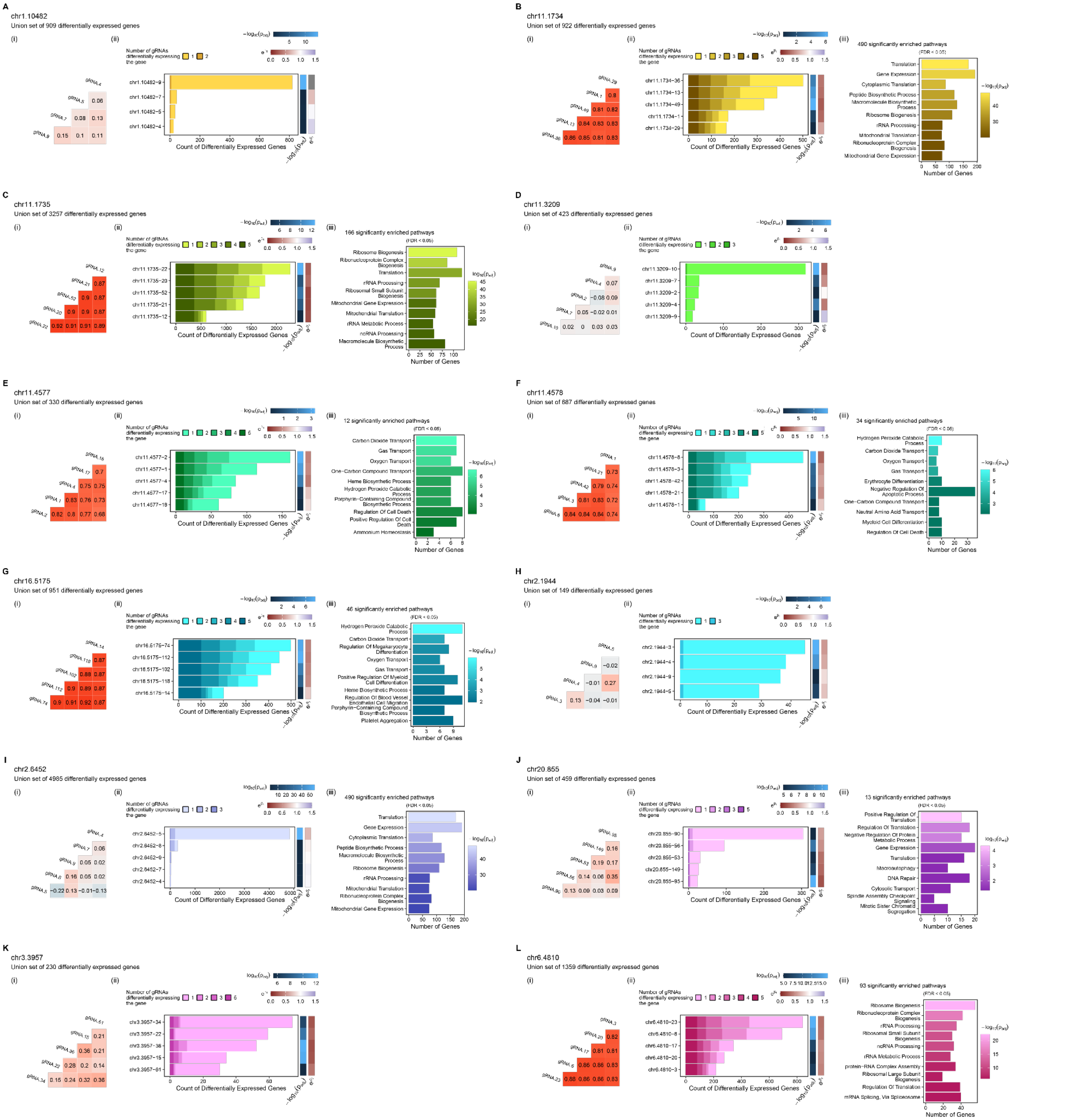


**Figure S21: Transcriptome-wide differential expression analysis of validation DHSs.** Each validation DHS is represented by three plots: **(i)** the transcriptome-wide correlation of the union set of differentially expressed genes across the DHS **(ii)** Counts of significantly differentially expressed genes for each gRNA in the DHS, partitioned by the number of gRNAs detected as affecting the gene's expression. Heatmaps on the right indicate magnitude of the gRNA's effect on the target gene ($e^{\beta_1}$ ) and the significance of that connection ($log_{10}\ p_{adj}$). **(iii)** Gene ontology analysis across the union set of differentially expressed genes for the gRNAs in the DHS.

**Figure S22: Replicate correlation of bulk 1-week screen raw gRNA counts.**

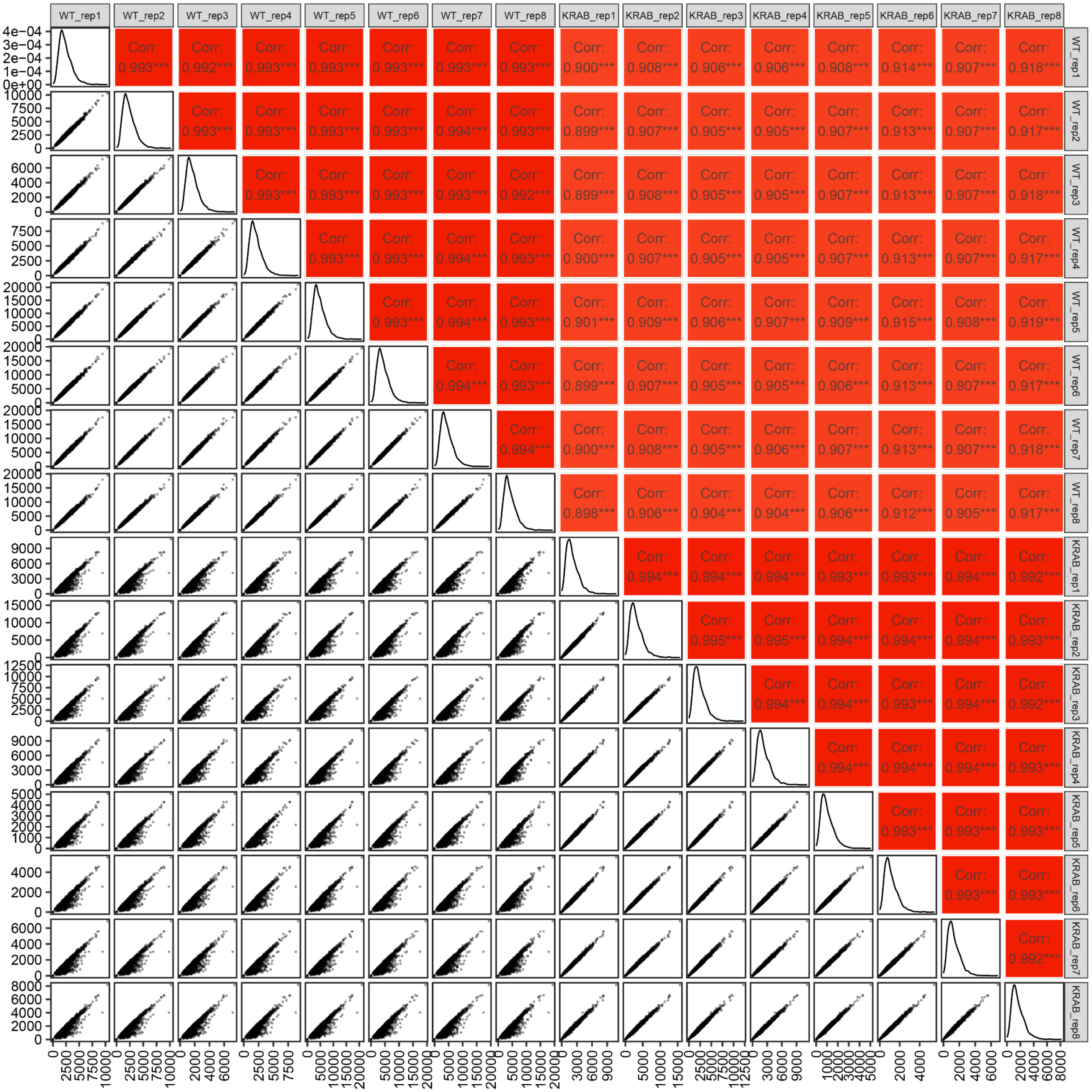


**Figure S22: Replicate correlation of bulk 1-week screen raw gRNA counts.** Correlation (Pearson's r) of raw gRNA sequencing read counts between replicates in the bulk 1-week screen in K562 cells. Each point represents a different protospacer sequence. Cells transfected with the same gRNA library used for the single cell screen into K562 cells with or without dCas9-KRAB. Cells without dCas9-KRAB are labelled WT. As expected, comparing WT to KRAB reveals protospacers that are depleted.

**Figure S23: DHS-level effects of 1-week cell fitness screen and comparison with validation screen.**

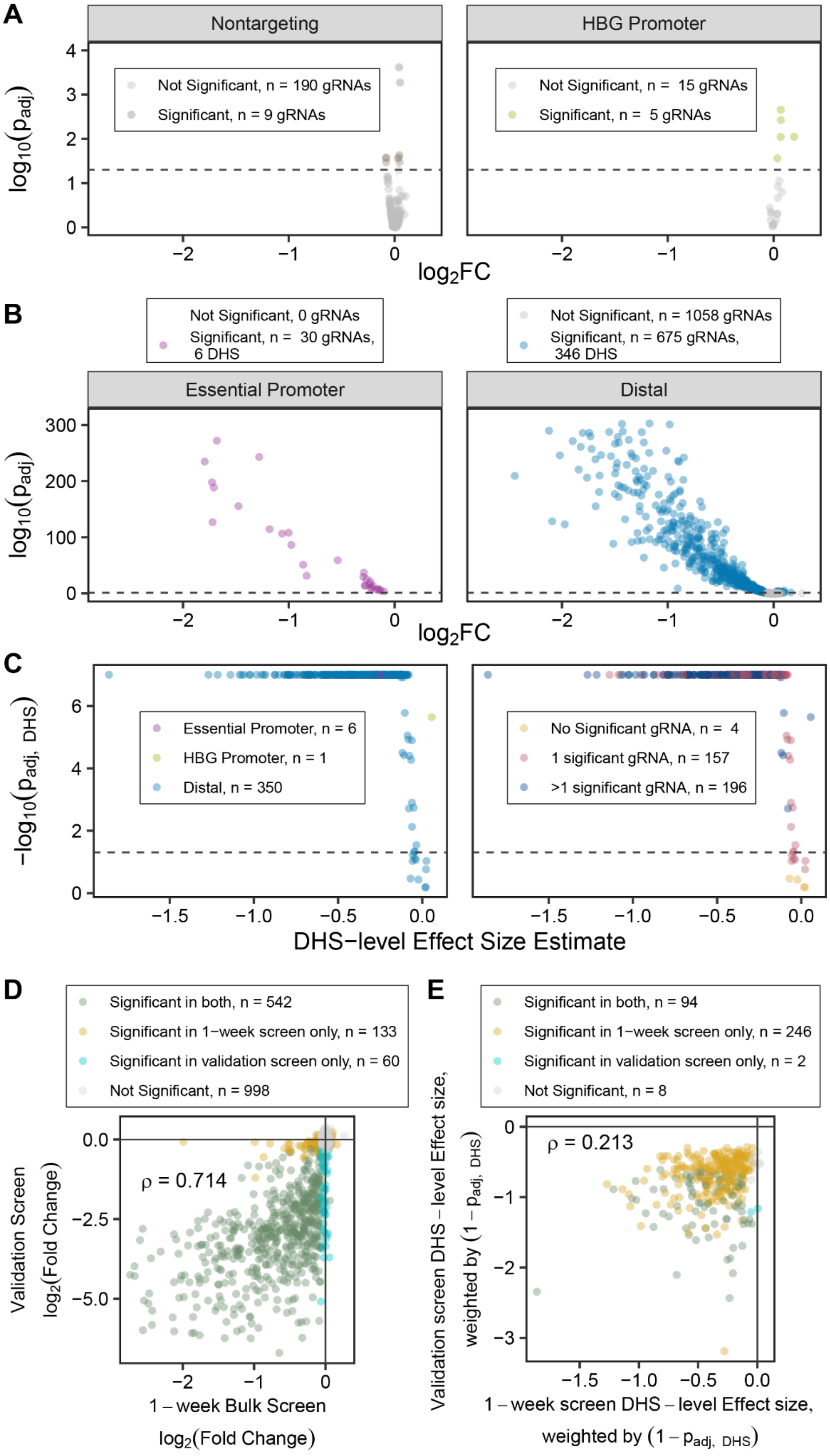


**Figure S23: DHS-level effects of 1-week cell fitness screen and comparison with validation screen. (A)** Fitness effects of nontargeting gRNAs and gRNAs targeting the HBG promoter. **(B)** Fitness effects of essential promoter controls and gRNAs targeting distal elements. **(C)** Aggregated DHS-level fitness effects and significances for the 1-week screen. Left: colored by DHS type. Right: colored by the number of individually significant gRNAs. **(D)** Correlation between gRNA-level fitness effect sizes in the validation and 1-week bulk screens. Spearman $\rho$ = 0.714, p < $10^{-16}$. **(E)** Correlation between DHS-level fitness effect sizes in the validation and 1-week bulk screens. Spearman $\rho$ = 0.213, p = 7.7x $10^{-5}$.

**Figure S24: Competition assay growth time courses of individual gRNAs, grouped by DHS.**

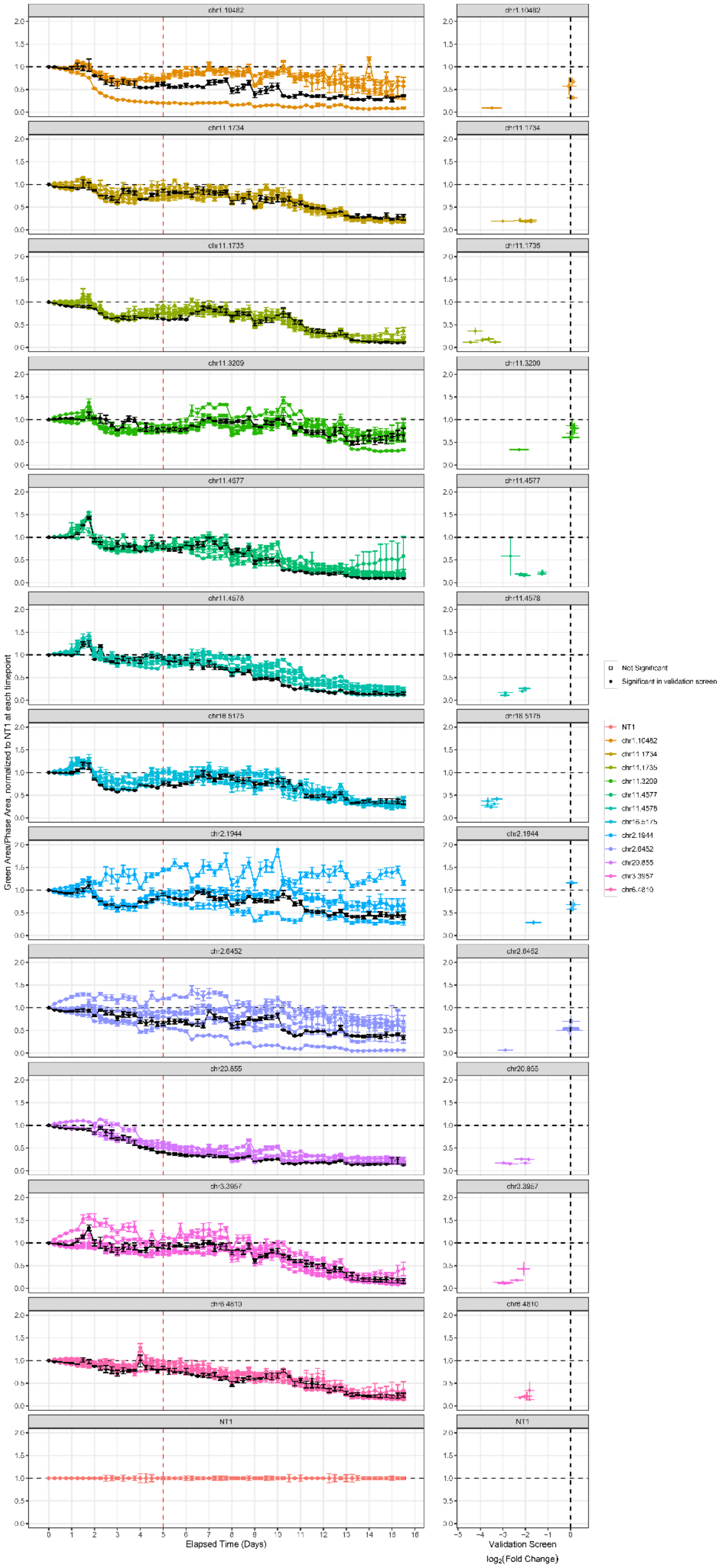


**Figure S24: Competition assay growth time courses of individual gRNAs, grouped by DHS.** Left column: Competition assay time courses of the ratio of green area to phase area, normalized to nontargeting signal at each timepoint. Each DHS is on a separate subplot; gRNAs are individual lines within the plot represented by 3-5 biological replicates with standard error bars. Right column: Correlation between validation screen ratio and the last time point of the competition assay. Error bars in both directions are standard error (3-5 biological replicates in y, 4 biological screen replicates in x).

**Figure S25: Distribution of discovery screen gRNA p-values by gRNA rank within the DHS.**

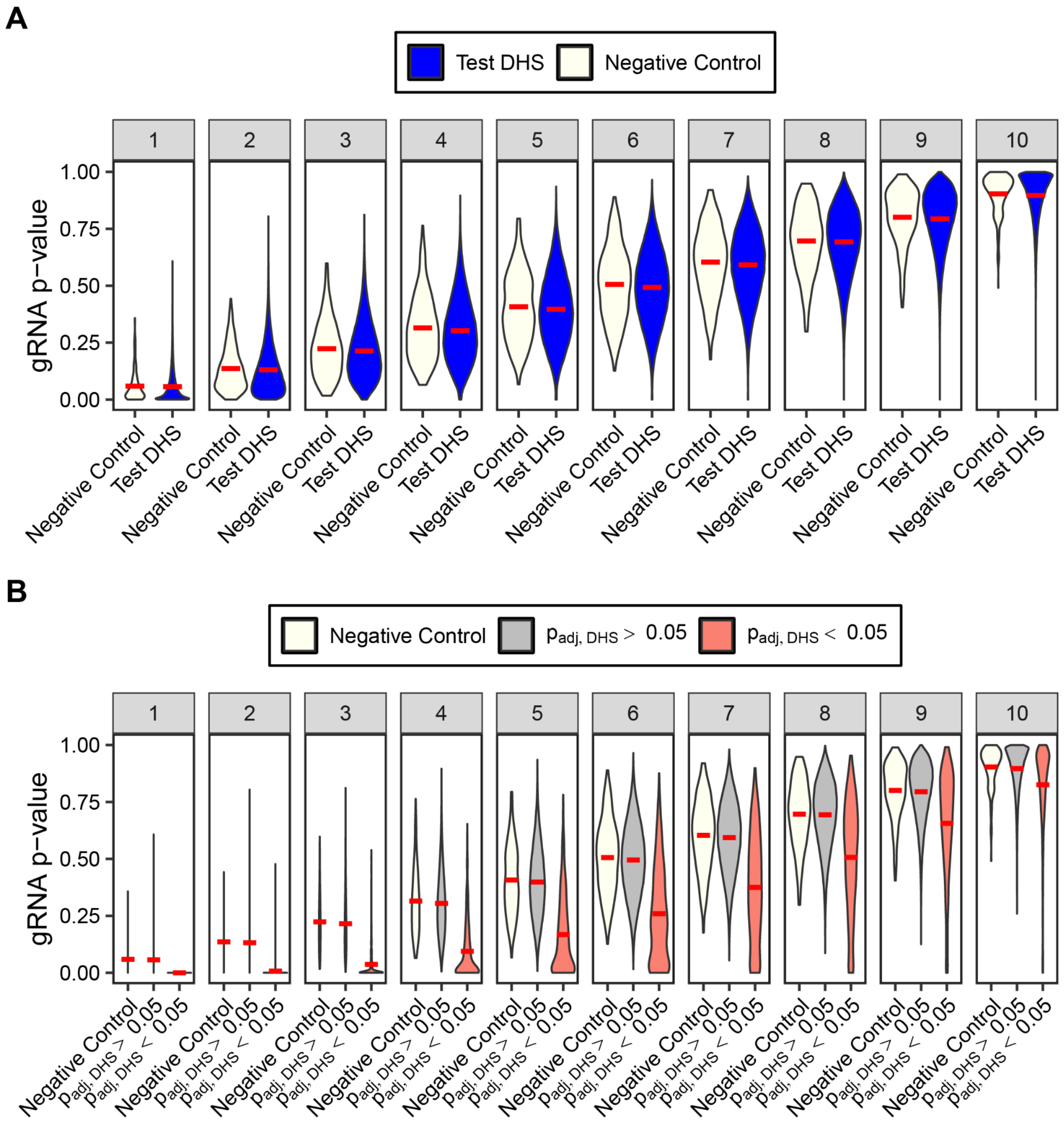


**Figure S25: Distribution of discovery screen gRNA p-values by gRNA rank within the DHS. (A)** Distribution of unadjusted gRNA p-values across order statistics (significance rank of gRNA within the DHS) in 2 distributions: 1) In white, the 282 negative control DHSs with exactly 10 gRNAs and 2) in blue, the 103,842 “test” DHSs in the discovery screen that have exactly 10 gRNAs. The red crossbar marks the mean of each p-value distribution. On average, there is no significant difference between the negative control and test DHSs p-values across all order statistics ($p>0.1$, Wilcox test). **(B)** The same distributions as in **(A)**, but with the “test” DHSs divided into significant and insignificant distributions. The red crossbar marks the mean of each distribution.
